# Diverse Pharmacogenomic Resistance Landscapes Across Epigenetic Drug Classes Revealed by Gastric Cancer Perturb-seq

**DOI:** 10.1101/2025.04.16.649236

**Authors:** Chang Xu, Haoran Ma, Taotao Sheng, Ming Hui Lee, Siti Aishah Binte Abdul Ghani, Laura Perlaza-Jimenez, Kie Kyon Huang, Shen Kiat Lim, Supriya Srivastava, Xuewen Ong, Su Ting Tay, Shamaine Wei Ting Ho, Zhi Xuan Ong, Angie Lay Keng Tan, Feng Zhu, Hassan Ashktorab, Alfred Sze-Lok Cheng, Anand D. Jeyasekharan, Shang Li, Ming Teh, Raghav Sundar, David R. Powell, Joseph Rosenbluh, Wei Peng Yong, Jimmy Bok-Yan So, Patrick Tan

## Abstract

Gene expression signatures (“molecular phenotypes”) are extensively utilized in cancer research. To study how gastric cancer (GC) molecular phenotypes are shaped by cell-intrinsic genetic alterations interacting with cell-extrinsic therapeutic pressures, we performed direct capture Perturb-seq (dcPerturb-seq) to interrogate >200 GC-related genes across 4 distinct epigenetic drug classes in multiple gastric lines. We captured 17.7 million pharmocogenomic expression interactions in 625,866 cells representing baseline and post-therapeutic molecular phenotypes. This single-cell pharmacogenomic compendium confirmed previously known gene-driven molecular phenotypes, elucidated poorly characterized genes, and uncovered novel gene dosage-molecular phenotype relationships. Molecular phenotypes in post-therapeutic surviving cells revealed diverse gene perturbation-associated pathways causing convergent drug resistance (EMT plasticity, cell cycle alterations, metabolic reprogramming), highlighting combinatorial strategies for restoring sensitivity. Mapping of *in vitro* molecular phenotypes to primary human GCs imparted prognostic information and insights into spatial heterogeneity. Comparative analysis of gene perturbations across therapies and lines revealed both conserved and context-specific molecular alterations. Our results illustrate how Perturb-seq approaches can systematically map diverse cancer-associated molecular phenotypes across multiple gene/drug/cell line interactions, yielding translational insights.

## Introduction

Cancer cells are dynamic cellular ecosystems exhibiting multiple functional phenotypes such as sustained proliferation, cell death resistance, and evasion of host immunity, ultimately culminating in malignancy [1]. Understanding how cancer phenotypes emerge from normal cells, through the complex interaction of germline and somatic genomic and epigenomic aberrations across decades of life, represents a cornerstone of molecular cancer research. In late-stage cancers, terminal phenotypes such as therapy resistance and metastasis represent key drivers of cancer mortality [2]. To capture the diverse functional states of cells in a structured format, transcriptome profiling has emerged as a powerful approach where high-dimensionality gene expression signatures can serve as “molecular surrogates” of functional phenotypes [3, 4]. Gene expression profiles (hereafter referred to as *molecular phenotypes*) have been used to identify clinically-relevant subtypes of cancer, predict patient prognosis, and to dissect disease mechanisms [5]. One important feature of molecular phenotypes is that they are often broadly “transportable” across different tumor types, and between pre-clinical models and *in vivo* settings [6]. Moreover, reflecting their biological coherence, molecular phenotypes associated with alterations in separate components of the same cellular process often exhibit high similarity [7], providing a basis for functional discovery.

The expression, composition, and coordination of molecular phenotypes is governed by many factors, both cell-intrinsic and extrinsic. Examples of cell-intrinsic factors driving specific molecular phenotypes include genomic alterations in classical oncogenes and tumor suppressor genes (eg *EGFR*, *TP53*, and *VHL* [8, 9]) and more recently chromatin modifier genes that are often mutated [10] and can elicit widespread epigenetic dysregulation [11]. Conversely, examples of cell-extrinsic factors linked to molecular phenotypes include nutrient availability [12], hypoxia [13], interactions with the tumor microenvironment [14], niche-specific cues [15], and therapeutic pressure from anti-cancer therapies [16–18]. Notably, relationships between molecular phenotypes and cell-intrinsic/cell-extrinsic factors are often complex and non-linear, being governed by gene-gene (GxG) and gene-drug (GxD) interactions and often further influenced by tissue lineage and micro-environmental context. For example, different molecular phenotypes have been shown to arise upon *KRAS* activation depending on the presence or absence of *TP53* function [19].

Recently, single-cell technologies for generating molecular phenotypes linked to gene perturbations have emerged [20]. Methodologies such as Perturb-seq and CROP-seq, which combine CRISPR genome editing with single-cell transcriptomics, have the potential for elucidating genetic perturbation-driven molecular phenotypes at scale [20–22]. These powerful approaches have been used to study differentiation trajectory impairments in TGF-β signalling [23] and to elucidate transcriptional regulation by mSWI/SNF complex subunits [24]. To date however, few studies have applied this approach to study molecular phenotypes associated with therapy resistance [25]. Here, we applied direct-capture Perturb-seq (dcPerturb-seq) in gastric cancer (GC), a leading cause of global cancer morbidity and mortality [26], exploring interactions between >200 cancer genes and four distinct epigenetic drugs in multiple lines. We selected therapies targeting epigenetic mechanisms due to the prominent role of epigenomics in regulating gene expression and cellular plasticity [27]. By mapping molecular phenotypes across 625,866 cells against 6 gastric cell lines, 226 genes, and 4 epigenetic drugs, we surveyed 4,746 gene/drug/cell line and 17.7 million expression interactions. Analysis of this single-cell pharmacogenomic compendium revealed dynamic interactions between genetic alterations, epigenetic therapies, and cellular responses, causing diverse molecular phenotypes underpinning gene-dosage effects, drug resistance, and context-specificities. Our work establishes a framework for a comprehensive atlas of drug-perturbation molecular phenotypes, yielding insights into how molecular phenotypes are modulated by context-specific drug action and cancer-associated genomic interactions.

## Results

### Direct Capture CRISPR Single-Cell Perturbation Sequencing (dcPerturb-seq) Platform

To systematically map molecular phenotypes associated with gene perturbations in GC, we compiled an initial list of 2,290 genes comprising known GC oncogenes, tumor suppressor genes (TSGs), cancer-related transcription factors (TFs), epigenetic regulators, and genes associated with GC progression (**Figure S1A**). For this proof-of-concept study, we filtered the list to 216 genes exhibiting either differential expression between primary GCs and matched normal tissues, relevance to GC supported by multiple studies, and detectible expression in gastric cell lines and single-cell RNA-seq epithelial profiles. We designed a Perturb-seq library containing four sgRNAs per target gene, and also included sgRNAs targeting essential *RPL* genes (positive controls) and non-targeting sgRNAs (negative controls). The total number of genes ultimately selected for screening was 226 (**Supplementary Table 1**).

Using direct capture Perturb-seq (dcPerturb-seq), we introduced the feature-barcoded lentiviral libraries into GES1 gastric epithelial cells (GES1, **Figure 1A**). Compared to standard Perturb-seq, dcPerturb-seq enables direct capture of sgRNAs, reduced lentiviral template switching, and is compatible with lineage tracing and molecular recording [28]. GES1 cells were chosen to function as a normal reference against which cancer lines could be compared [29] (**Supplementary Table 2** lists dcPerturb-seq gene protein-coding mutations across all lines). For GES1, we recovered 74,728 high-quality cells of which 32,283 (43%) were successfully annotated with CRISPR sgRNAs. Of these, 28,721 cells (89%) carried a single sgRNA across the 998 sgRNAs (**Figure S1B)**. We performed stringent quality control (QC) analyses to ensure data quality. First, we confirmed the expected functional impact of the dcPerturb-seq library by observing significant reductions in cell populations containing sgRNAs targeting *RPL* essential genes (**Figure S1C;** p=0.0097) and DepMap [30]-predicted essential genes such as *CDK1*, *TOP2A* and *AURKB* (**Figure S1D** and **S1E;** overall p=1.7×10^-6^). Second, we performed single cell long-read RNA-seq to verify accurate CRISPR editing, and observed an average of 4.21-fold higher indel deletion rates in cells transduced with targeting sgRNAs (passing read depth thresholds) compared to non-targeting control cells (**Methods**). Third, we assessed impact of the sgRNAs on target gene expression levels. Here, we caveat that as CRISPR perturbations target genomic DNA (unlike siRNA and shRNA perturbations which directly target RNA), detecting CRISPR-associated effects on target gene expression levels are often reliant on multiple factors, such as baseline target gene expression levels, sufficiency of cell numbers, and the secondary effects of nonsense-mediated decay (NMD) which has variable efficiency [22, 24] (**Figure S1F**). Nevertheless, for >70% of targets we observed significantly reduced expression of target genes compared to cells with sgRNAs targeting other genes or control cells (p value < 2.2 × 10^-16^, binomial test) (**Figure S1G** and **S1H**). Consistent with CRISPR perturbations exerting their functional impact primarily at the genomic (DNA) and not transcriptomic (RNA) level, no significant correlations were detected between the extent of target expression reduction and phenotypic effects on cell survival (**Figure S1I**).

**Figure 1.**
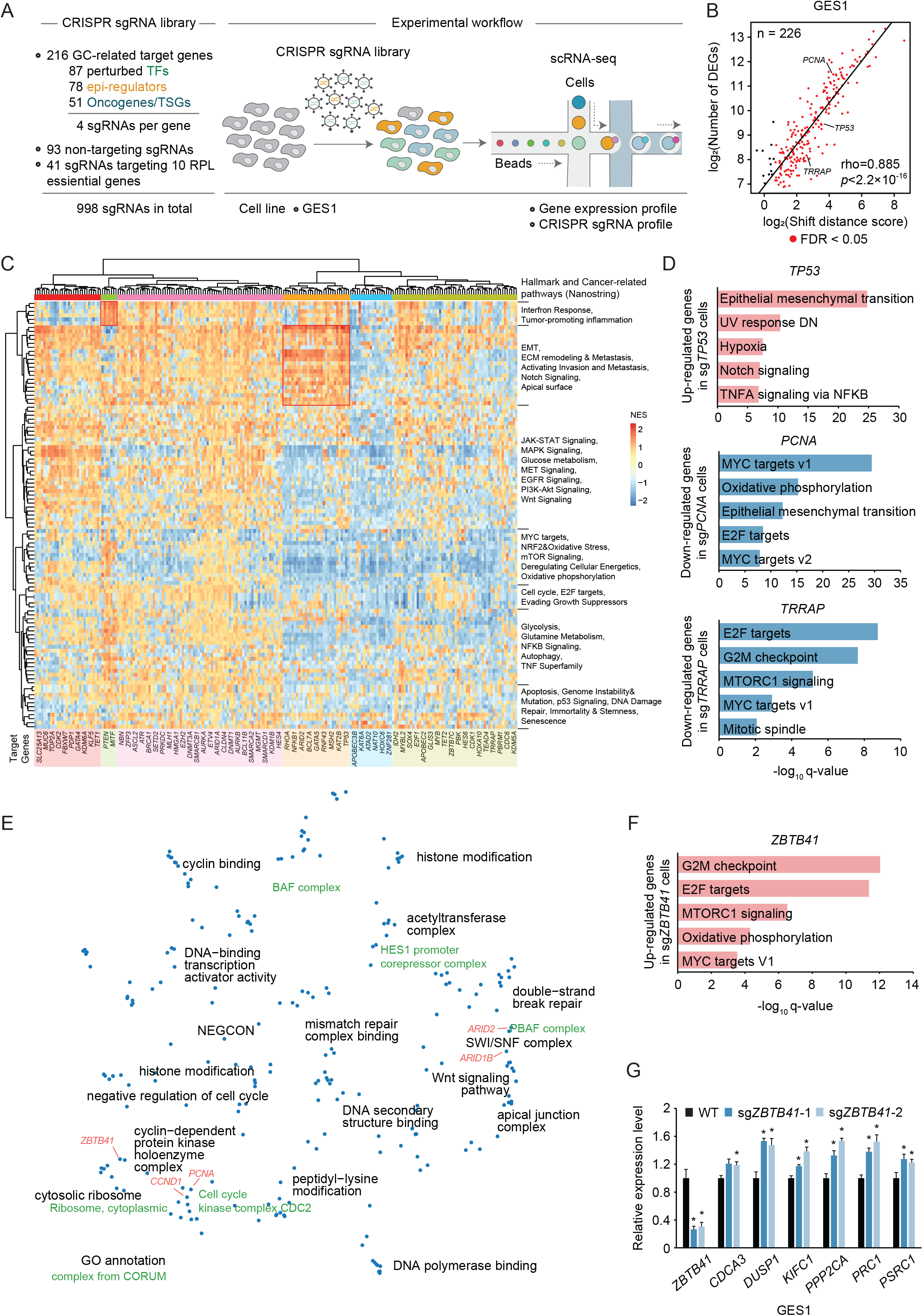
Annotating Gene Functions Using dcPerturb-seq. (A) Schematic representation of dcPerturb-seq experimental design and workflow. (B) Correlation between shift distance scores (defined by Hotelling’s *T*^2^ test) and numbers of differentially expressed genes (DEGs) for each perturbed target gene in GES1 cells. Perturbations were compared against wild-type (WT) cells. Significance was determined using Pearson correlations. (C) Heatmap showing normalized enrichment scores (NES) for pathway analysis (rows) and hierarchical clustering of perturbation-associated molecular phenotypes (rows). Gene names at the bottom represent selected genes from the dcPeturb-seq panel (columns). (D) Enrichment analysis of biological pathways of perturbed genes (*TP53*, *PCNA*, and *TRRAP)* in GES1 cells. x-axis values represent log_10_(q-value), where q-values are Benjamini-Hochberg (BH)-adjusted p-values from the pathway enrichment test. (E) Two-dimensional Minimum-Distortion Embedding (MDE) plot where each dot represents a gene perturbation, positioned by the mean embedding across all cells for each perturbation. Clusters were annotated using Gene Ontology (GO) pathway enrichments (black) and CORUM database annotations (green). (F) Biological pathway enrichment analysis of molecular phenotypes associated with *ZBTB41* gene perturbation in GES1 cells. Values on the x-axis represent log_10_(q-value), where q-values are BH-adjusted p-values from the pathway enrichment test. (G) qRT-PCR results showing relative expression levels of *ZBTB41*, *CDCA3*, *DUSP1*, *KIFC1*, *PPP2CA*, *PRC1* and *PSRC1* genes in WT and *ZBTB41*-perturbed GES1 cells (n=3; mean ± SD) (*, p < 0.05, two-sided *t*-test).

To assess molecular phenotypes linked to specific gene perturbations, transcriptomic profiles of cells with sgRNAs targeting the same gene were aggregated and normalized (**Methods**). We quantified global shifts in mean expression profiles (termed “shift distance scores”) using Hotelling’s *T*^2^ test [31] between cells with target-specific sgRNAs against non-targeting control cells (**Methods**). To identify specific differentially expressed genes (DEGs), we applied an Anderson-Darling (AD) test [7], known for its sensitivity to detect transcriptional changes in subsets of cells. We observed positive correlations between the magnitude of global transcriptional shifts and the numbers of DEGs (average 127 cells/gene for GES1) (**Figure 1B**) and confirmed the robustness of this approach by repeating the analyses after down-sampling 80% and 60% of cells, observing consistent results (**Figure S2A**). We validated our methodology by applying TRADE, a recently published single-cell differential expression algorithm [32] (**Figure S2B**), confirming significant positive correlations between DEG numbers/shift distance scores identified by our method and genome-wide transcriptome-wide impact (TI) levels computed by TRADE (**Figure S2C**). Further, as an additional QC and reflecting our dataset’s biological coherence, sgRNAs targeting the same gene exhibited significant positive correlations in TI levels (F value = 1.619, p value = 0.041, one-way ANOVA test), and target genes previously identified as evolutionarily constrained and loss-of-function intolerant (LOEUF) were associated with higher TI scores (**Figure S2D**) [33].

### dcPerturb-seq reveals known and novel functional gene annotations

Unsupervised clustering of the 216 perturbation-associated molecular phenotypes revealed several clusters (**Figure 1C**). To validate individual perturbation-associated molecular phenotypes, we evaluated candidate cancer-related genes. In sg*TP53* targeting cells, we observed upregulation of DEG signatures related to epithelial mesenchymal transition (EMT) pathways, while cell cycle related signatures were downregulated in cells with sg*PCNA* and sg*TRRAP* perturbations, consistent with previous reports [34–36] (**Figure 1D**). We also observed similarities in molecular phenotypes for perturbed targets within the same protein complex. For example, the cell cycle kinase complex members *PCNA* and *CCND1* exhibited similar transcriptional signatures, as did *BRCA1* and *BRCA2* within the BRCC complex (**Figure S2E**). Pathway enrichment analysis identified a set of gene perturbations associated with up-regulation of inflammatory related processes - this cluster included *PTEN*, the deletion of which has been previously reported to increase NF-κB activity [37]. In another cluster, EMT, metastasis, and Notch Signaling pathways were activated in cells with perturbed EMT-related genes (eg *RHOA*, *ARID2* and *TP53*).

To visualize these perturbation associations beyond pair-wise gene-gene relationships, we performed dimensional reduction of the perturbation-associated molecular phenotypes using a machine-learning-based Minimum-Distortion Embedding (MDE) solver [38], and projected the embeddings into a 2D UMAP (Uniform Manifold Approximation and Projection) followed by unsupervised clustering (see **Methods**). We identified 21 discrete clusters and annotated each cluster using Gene Ontology (GO) pathways and CORUM, a database of experimentally validated protein-protein interactions [39] (**Figure 1E**). The 21 clusters showed a clear distribution of discrete biological processes including histone modification, double-strand break repair, apical junction complex and cyclin-dependent protein kinases. Genes associated with similar protein complexes clustered in close proximity such as *ARID2*/*ARID1B* in the PBAF complex and *CCND1*/*PCNA* in the cell cycle kinase complex.

To ask if leveraging these perturbation clusters might facilitate the annotation of less characterized genes, we considered *ZBTB41* (Zinc Finger And BTB Domain Containing 41), a poorly annotated gene in the literature (7 papers in Pubmed as of Oct 2025). In UMAP space, *ZBTB41* clustered with cell cycle related genes/complexes, suggesting a role in cell cycle regulation. Pathway analysis of *ZBTB41* perturbation-associated molecular phenotypes confirmed significant enrichment of cell cycle related processes including G2M checkpoint regulation, mitotic cell cycle and nuclear division processes in GES1 cells (**Figure 1F**). To validate these findings, we independently perturbed *ZBTB41* in GES1 and HFE145 normal gastric epithelial cells and again observed up-regulation of cell cycle related targets (eg *CDCA3*, *DUSP1* and *KIFC1*) (**Figure 1G** and **Figure S3A**). Functional cell cycle analysis by flow cytometry confirmed that *ZBTB41* perturbations resulted in a shift of cell cycle distributions, marked by a decreased proportion of G1-phase cells and an increased proportion of S-phase cells without fitness costs (**Figure S3B-F**). In summary, these data demonstrate the efficacy of the dcPerturb-seq platform in simultaneously capturing molecular phenotypes of multiple target genes after perturbation, which can facilitate functional assignments for poorly annotated genes.

### Exploring Effects of Expression Dosage on Molecular Phenotypes

Little is known about how linear alterations in target gene mRNA levels (“expression dosage”) can influence the expression of downstream transcriptional signatures [40, 41]. We explored if dcPerturb-seq might allow us to evaluate how molecular phenotypes are influenced by the expression dosage of target genes at the single-cell level. For this analysis, we *a priori* defined three potential classes of perturbation targets - Dosage-sensitive (DS), Dosage-Linear (DL), and Dosage Tolerant (DT). DS genes correspond to targets where modest changes in gene expression can already cause transcriptional effects analogous to complete expression depletion, akin to genetic haploinsufficiency [42]. DL genes correspond to genes where reductions in expression are associated with correspondingly graded alterations in transcriptional phenotype. Lastly, DT genes represent targets showing transcriptional changes only when target expression is severely reduced (**Figure 2A**).

**Figure 2.**
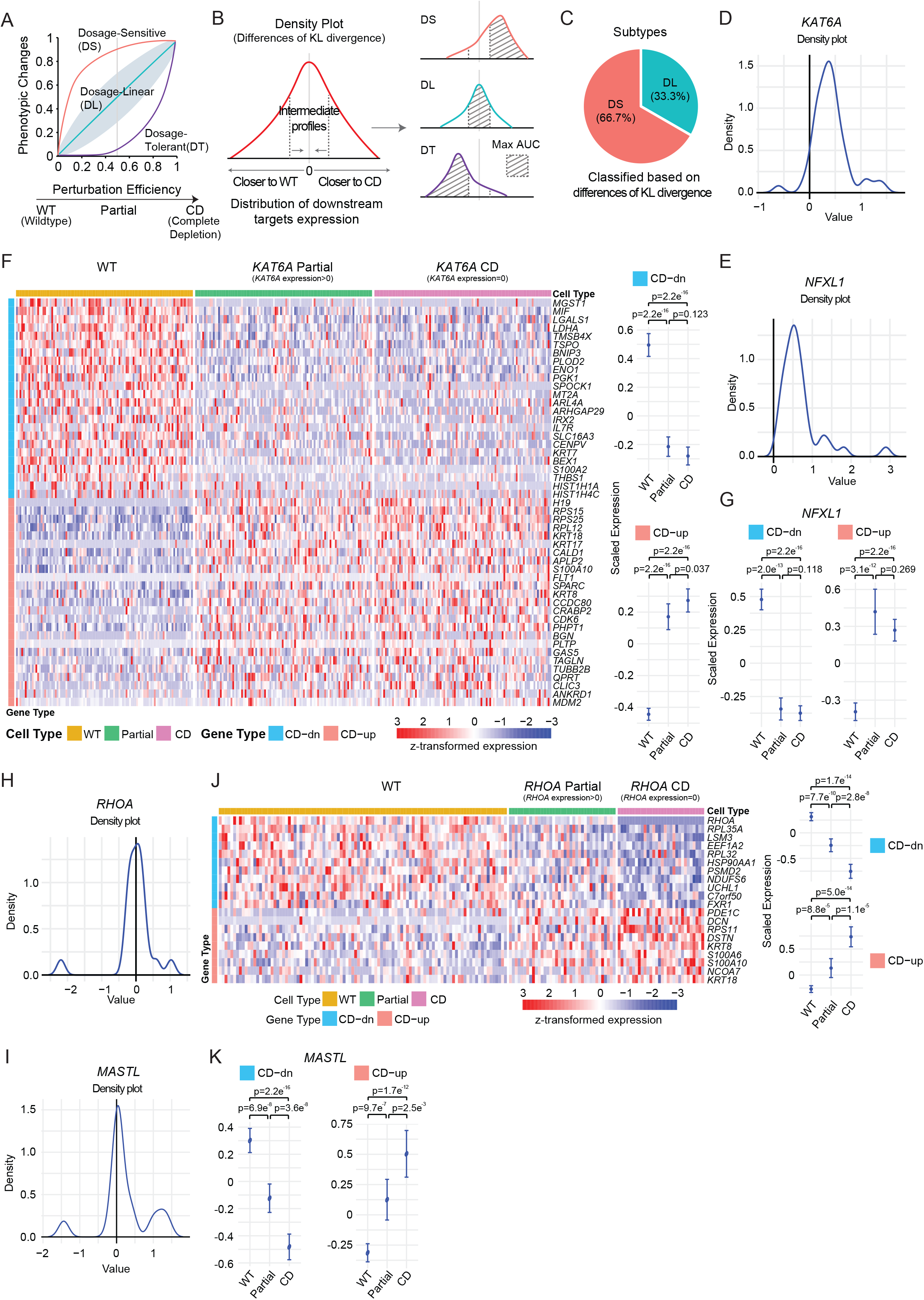
Target Gene Expression Dosage and Molecular Phenotypes. (A) Diagram showing alternative classes of transcriptional responses to gene perturbations. Perturbation efficiency (x-axis) indicates the degree of reduction in target gene expression. Phenotypic changes (y-axis) represent the scale of consequent transcriptional response. (B) Overview of classification methods used to define categories of transcriptional responses associated with Figure 2A. (C) Distribution of target gene expression dosage against transcriptional response subtypes, determined using Kullback-Leibler (KL) divergence. (D) Density plot showing shifts in cell profiles with partial *KAT6A* expression depletion (Partial) compared to profiles observed in cells with complete *KAT6A* expression deletion (CD). Values on the x-axis represent differences between the KL-defined distances of (*KAT6A* Partial, WT) and distances of (*KAT6A* Partial, *KAT6A* CD). (E) Density plot showing shifts in cell profiles with partial *NFXL1* expression depletion compared to profiles observed in cells with complete *NFXL1* expression deletion. (F) Heatmap of downstream transcriptional responses in WT, *KAT6A* partially-depleted (Partial; *KAT6A* expression>0) and *KAT6A* completely-depleted (CD; *KAT6A* expression=0) cells (left), along with a summary of scaled expression (right) (*P* values calculated using Wilcoxon rank sum test). Columns represent cells belonging to either WT, *KAT6A* Partial or *KAT6A* CD groups. Rows represent genes differentially expressed between *KAT6A* CD and WT cells, identified by the Wilcoxon rank sum test. Row colors denote the direction of regulation in the *KAT6A* CD group relative to WT cells (blue, downregulated; orange, upregulated). (G) Summary of scaled downstream transcriptional responses in WT, *NFXL1* partially-depleted and *NFXL1* completely-depleted cells (*P* values calculated using Wilcoxon rank sum test). (H) Density plot showing intermediate molecular phenotype expression in cells with partial *RHOA* expression depletion compared to WT cells and cells with complete *RHOA* expression deletion. (I) Density plot showing intermediate molecular phenotype expression in of cells with partial *MASTL* expression compared to WT cells and cells with complete *MASTL* expression depletion. (J) Heatmap of downstream transcriptional responses in WT, *RHOA* partially-depleted (Partial) and *RHOA* completely-depleted (CD) cells (left), with a summary of scaled expression (right) (*P* values calculated using Wilcoxon rank sum test). Columns represent cells belonging to either WT, *RHOA* Partial or *RHOA* CD groups. Rows represent genes differentially expressed between *RHOA* CD and WT cells, identified by the Wilcoxon rank sum test. Row colors denote the direction of regulation in the *RHOA* CD group relative to WT cells (blue, downregulated; orange, upregulated). (K) Summary of scaled downstream transcriptional responses in WT, *MASTL* partially-depleted and *MASTL* completely-depleted cells (*P* values calculated using Wilcoxon rank sum test).

For each of the 216 target genes, we stratified cells into discrete populations showing either partial or complete depletion of target expression, conducting differential analysis against Wildtype (WT) negative control cells. To ensure data robustness, we applied a stringent single cell-based differential analysis pipeline focusing on strong and significant downstream alterations (**Methods**). We developed a Kullback–Leibler (KL) divergence-based method to classify Perturbation-Phenotype patterns and employed for validation a distinct additional Euclidian distance-based approach (**Figure 2B**) (**Methods**). Under this analysis framework, two-thirds of dosage-effect patterns fell into the DS category (66.7%), while the remainder belonged to the DL category (33.3%) (**Figure 2C**). No DT patterns were detected among the 216 perturbed targets, potentially due to limited cell numbers or initial gene selection (see **Discussion**). This finding suggests that for many of the genes selected in this study, modest changes in expression may be sufficient to generate robust molecular phenotypes.

As examples of DS genes, cells with partial *KAT6A* and *NFXL1* depletion (target expression >0) exhibited near-complete global shifts toward profiles seen in completely depleted cells (target expression = 0) on density plots (**Figure 2D-E**). Examining gene signature compositions, we confirmed that cells with partial *KAT6A* and *NFXL1* depletion displayed molecular phenotypes closely resembling those of completely depleted cells (**Figure 2F-G** and **Figure S4A**). This finding is consistent with previous studies reporting *KAT6A* as a haploinsufficient gene [43]. For DL genes such as *RHOA* and *MASTL*, density plots indicated an intermediate state between WT control and CD cells (**Figure 2H-I**), and expression profiles of partially-depleted cells exhibited intermediate levels of enrichment between WT negative control and completely depleted cells (**Figure 2J-K** and **Figure S4B**). These results suggest that dcPerturb-seq can provide insights into the sensitivity of perturbation-associated molecular phenotypes to expression dosage, which would be challenging to discern by bulk RNA analysis as the latter aggregates average expression across a large cell population.

### dcPerturb-seq Perturbations Reveal Distinct Molecular Phenotypes Associated with Epigenetic Pharmacology

To explore how dcPerturb-seq gene perturbations might influence the sensitivity or resistance of cells to therapeutic pressure, we focused on epigenetic drugs due to their ability to elicit widespread transcriptional re-programming [44]. After two weeks of library infection, independent pools of GES1 cells were treated with representative drugs covering four distinct epigenetic pathways - histone deacetylase (HDAC) inhibition (SAHA) [45]; DNA methyltransferse (DMNT) inhibition (5-azacytidine; 5-AZA) [46]; Bromodomain and Extraterminal Domain Inhibition (BETi) (JQ1) [47]; and Polycomb repressive complex 2 (PRC2) inhibition (GSK126) [48]. Drug exposures were maintained for another two weeks at concentrations causing 40-60% cell death. In total, we generated 121,672 high-quality cells across five experimental conditions: DMSO/Control (28,721 cells), SAHA (41,123 cells), 5-AZA (11,432 cells), JQ1 (29,736 cells), and GSK126 (10,660 cells) (**Figure 3A**).

**Figure 3.**
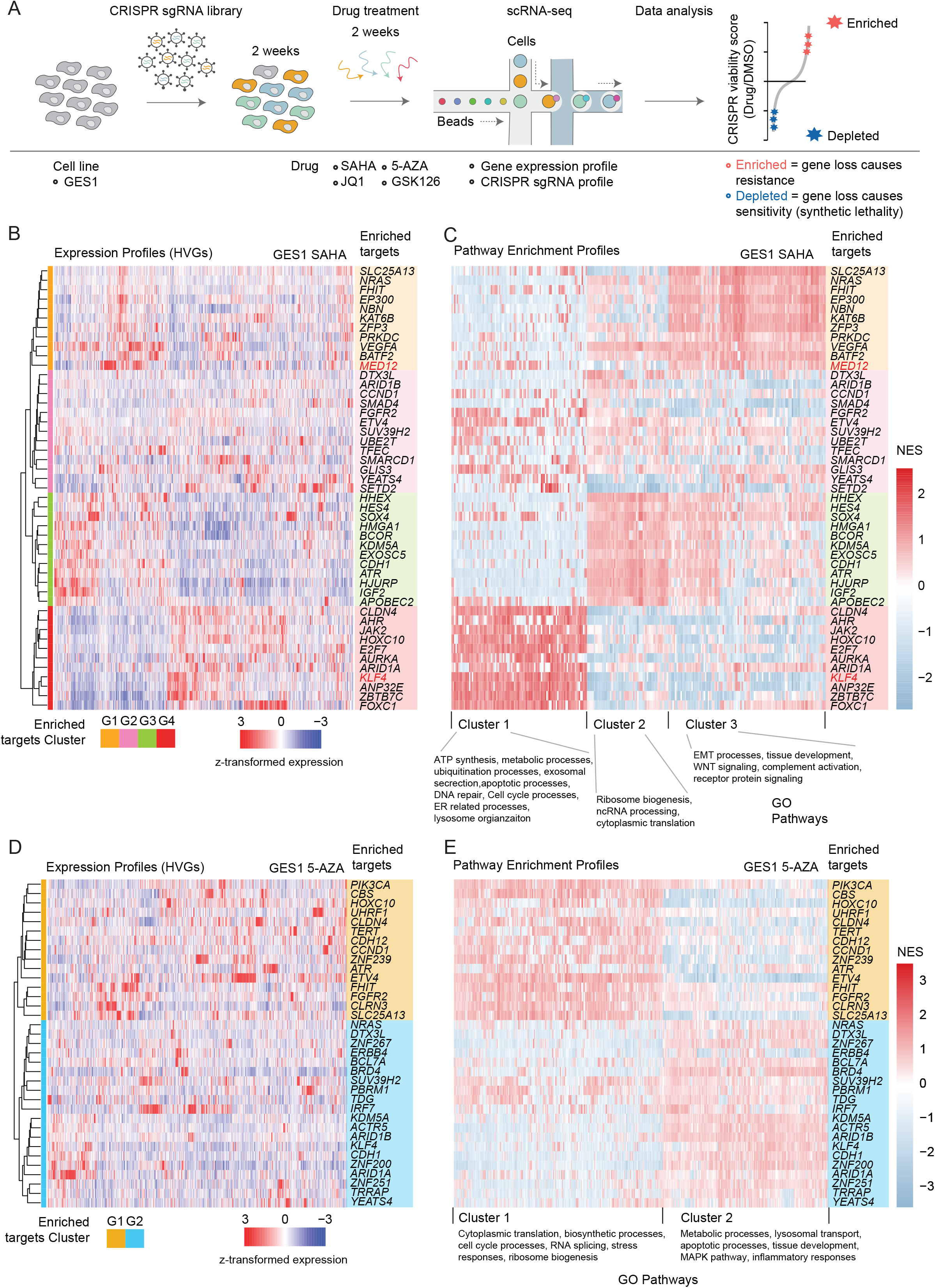
dcPerturb-seq Molecular Phenotypes Reveal Distinct Pathways Associated with Therapy Resistance. (A) Schematic of dcPerturb-seq platform combined with epigenetic drug treatments (SAHA, 5-AZA, JQ1 and GSK126). Drugs were added after two weeks of dcPerturb-seq library transfection. (B) Hierarchical clustering of enriched gene perturbation targets in response to SAHA treatment, based on *Z* transformed expression profiles. Rows represent individual SAHA-enriched target gene perturbations; columns represent highly variable genes (HVGs) defined across the SAHA-enriched targets, at the level of dcPerturb-seq molecular phenotypes. (C) Heatmap depicting NES scores for Gene Ontology (GO) pathways associated with enriched gene perturbation targets in response to SAHA treatment. Rows represent individual SAHA-enriched target gene perturbations in the same order and cluster as in Figure 3B; Columns represent GO pathways. Pathways were clustered by NES values and annotated by their cellular functions. (D) Hierarchical clustering of enriched gene perturbation targets in response to 5-AZA treatment, based on *Z* transformed expression profiles. Rows represent individual 5-AZA-enriched target gene perturbations; columns represented highly variable genes (HVGs) defined across the 5-AZA-enriched targets, at the level of dcPerturb-seq molecular phenotypes. (E) Heatmap depicting NES scores for Gene Ontology (GO) pathways associated with enriched gene perturbation targets in response to 5-AZA treatment. Rows represent individual 5-AZA-enriched target gene perturbations in the same order and cluster as in Figure 3D; Columns represent GO pathways. Pathways were clustered by NES values and annotated by their cellular functions.

Cells were first analyzed for CRISPR sgRNA counts (“viability scores”). Viability scores based on single-cell measurements correlated highly with pseudo-bulk CRISPR read-based viability measurements (**Figure S5A**, **Supplementary Table 3** provides CRISPR viability scores before and after normalization to DMSO conditions). We found that ∼20% and ∼15% gene perturbations were associated with either heightened resistance or sensitivity to at least one epigenetic therapy [49]. To validate the viability scores, we examined an independent genome-wide CRISPR screening dataset of JQ1-treated MCF7 breast cancer cells [50] and observed a significantly correlated pattern of gene enrichment and depletion to our viability scores (p<0.05; **Figure S5B**). To orthogonally validate the gene perturbation/therapy response interactions, we also independently perturbed three candidate targets (*MED12*, *KLF4* and *EP300*) in GES1 cells. The independently perturbed cells all exhibited resistance phenotypes to SAHA treatment, displaying increased cell viability compared to control (WT) cells after SAHA exposure (**Figure S5C**).

Therapy resistance is a downstream functional hallmark that can arise from different upstream mechanisms [16, 51]. We asked if dcPerturb-seq molecular phenotypes might provide insights into the heterogeneity of pathways leading to epigenetic drug resistance. As an initial use-case, we focused on SAHA and the top 50 resistance-associated gene targets exhibiting post-SAHA viability score enrichment (“enriched targets”) (**Figure S6A**). Unsupervised clustering of post-SAHA molecular phenotypes revealed four distinct clusters (G1-G4) (**Figure 3B**), with less distinct clustering patterns observed for non-enriched targets (**Figure S6B**) (**Methods**). We performed GSEA analysis to elucidate pathways associated with the G1-G4 clusters (**Figure 3C**; see **Methods**). Pathways associated with EMT and WNT signaling were significantly up-regulated in Cluster G1, ribosome biogenesis in Cluster G3, and Cluster G4 upregulated pathways involved in ATP synthesis, metabolism, cell cycle, and endoplasmic reticulum activity. Cluster G2 exhibited minimal cluster-specific pathway alterations. Focusing on specific genes within each cluster, we selected *MED12* in G1 and *KLF4* in G4 for closer inspection. dcPerturb-seq of *MED12* in Cluster G1 caused up-regulation of cell-substrate adhesion pathways after SAHA treatment **(Figure S6C)**, while perturbation of *KLF4* in Cluster G4 caused an enrichment of cell cycle-related processes post SAHA challenge (**Figure S6D**). In independent experiments, we validated the up-regulation of cell substrate adhesion pathway-related targets at the bulk mRNA level in *MED12* perturbed cells treated with SAHA compared to DMSO (**Figure S6E**-**F**), and up-regulation of cell cycle-related genes (*ACTL6A*, *MKI67*, *KIF18A*) in SAHA-treated *KLF4*-perturbed cells (**Figure S6G-H**). These results showcase how dcPerturb-seq can uncover heterogenous molecular phenotypes converging onto a common drug-resistant functional cancer hallmark.

We applied a similar analysis framework to 5-AZA, JQ1 and GSK126. For 5-AZA, we identified two post-Aza transcriptional clusters associated with 35 enriched target genes, with one cluster displaying up-regulation of processes related to cytoplasmic translation, cell cycle and stress response, while the other cluster was enriched for metabolic activity, lysosomal transport and MAPK pathways (**Figure 3D-E**). One of these clusters contained *ARID1A* and *ARID1B*, both of which have been reported to exhibit increased expression in cell lines after DNMT inhibitor treatment [52]. For JQ1, we identified three post-JQ1 transcriptional clusters associated with 41 enriched targets (**Figure S7A-B**). Of these, cluster 3 exhibited up-regulation of histone deubiquitination processes and contained *FBXW7*, the loss of which has been reported to mediate BRAF degradation and resistance to BET inhibitors in leukemia [53]. Lastly, for GSK126, we detected a post-GSK126 resistance signature associated with ARID family (*ARID1B* and *ARID2*) perturbed cells, characterized by up-regulation of processes related to spliceosomal assembly, cytoplasmic translation and ribosome biogenesis (**Figure S7C-D**). This transcriptional signature was distinct from that observed in JQ1-treated *ARID* perturbed cells, indicating that distinct molecular phenotypes can arise even to the same genetic perturbation, under different extrinsic drug treatments.

We evaluated the extent to which dcPerturb-seq resistance-associated molecular phenotypes identified under one treatment might be similarly enriched across other treatments. Specifically, we compared molecular phenotypes associated with SAHA resistance to the molecular phenotypes of enriched targets under 5-AZA, JQ1, and GSK126 treatment (**Figure S7E)**. Suggesting the potential existence of conserved resistance mechanisms, molecular phenotypes related to SAHA gene cluster G1 and G4 were also enriched in resistance-associated phenotypes for JQ1 and GSK126 (p < 0.05). A minor enrichment of SAHA gene cluster G1-related phenotypes was also observed for 5-AZA (p=0.05).

### Translational Application of dcPerturb-seq Molecular Signatures

To evaluate the potential translational relevance of the dcPerturb-seq molecular signatures, we focused on SAHA-enriched targets (**Figure 4A**). First, we benchmarked our SAHA-enriched targets against the existing literature for previously-reported roles in therapy resistance, assessing both causal and correlative evidence (**Supplementary Table 4**). We confirmed several targets previously associated with therapeutic resistance in other cancers including *EP300* [54] and *SMAD4* [55], and uncovered new candidates, such as *HHEX* and *GLIS3*. Next, we hypothesized that collateral molecular vulnerabilities conferred by the upregulation of these resistance mechanisms might highlight strategies to overcome drug resistance. Specifically, we selected representative perturbation targets from SAHA-enriched clusters: *EP300* (G1) and *ARID1A* (G4) (**Figure S8A**). Cells harbouring these gene perturbations were treated with a combination of SAHA and specific inhibitors targeting their predicted resistance pathways. We found that *EP300*-perturbed cells under SAHA challenge exhibited increased sensitivity to inhibitors targeting EMT and WNT-related pathways, but not *ARID1A*-perturbed cells. Conversely, *ARID1A*-perturbed cells treated with SAHA showed heightened sensitivity to cell cycle inhibition, but not *EP300*-perturbed cells (**Figure 4B** and **Figure S8B**). These findings demonstrate potential strategies for overcoming drug resistance, by forecasting and targeting resistance mechanisms associated with specific gene perturbations.

**Figure 4.**
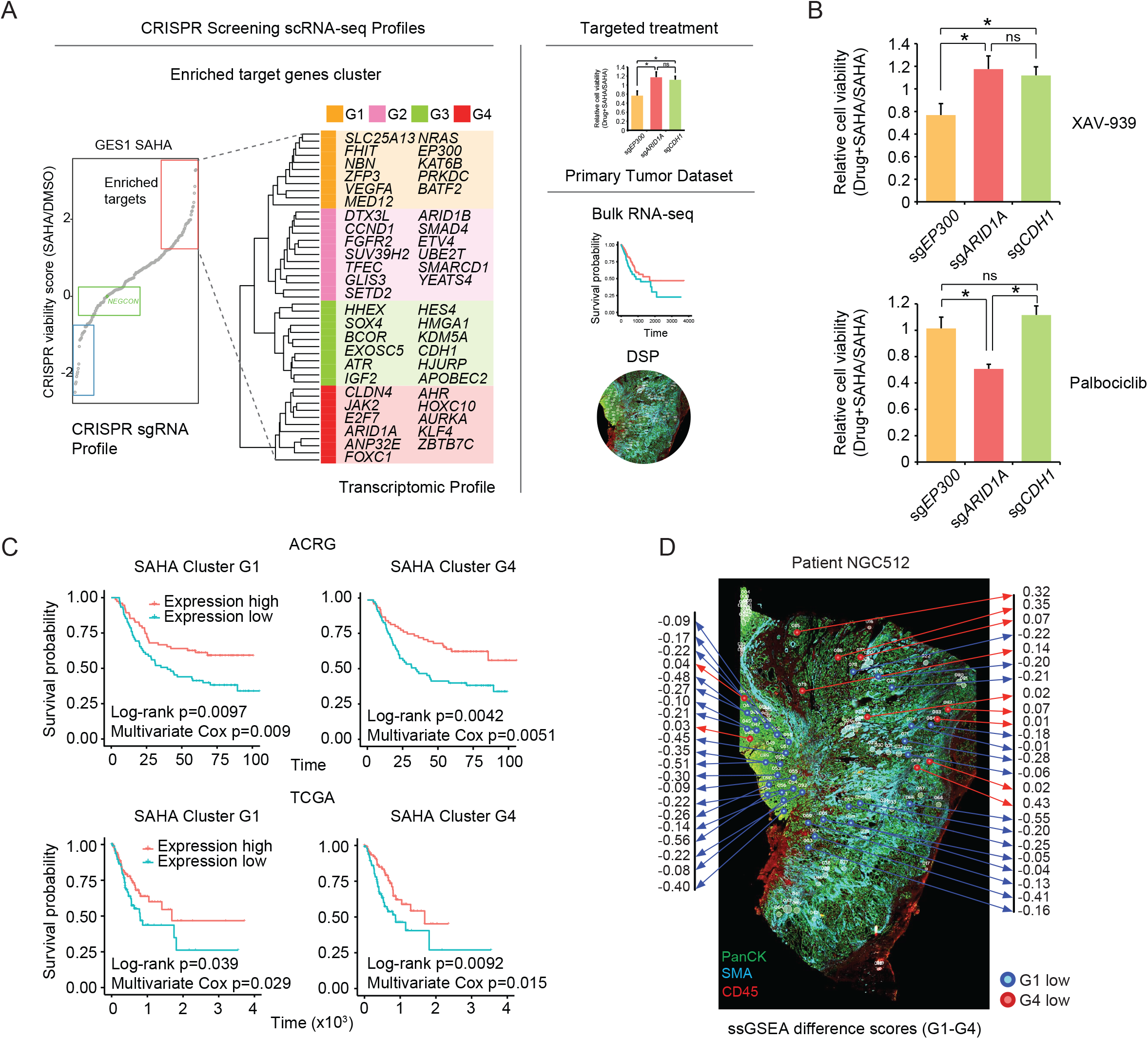
Translational Application of dcPerturb-seq Molecular Phenotypes. (A) Diagram illustrating the application of molecular phenotypes associated with enriched target genes (left) to targeted treatment, primary tumor expression data and NanoString GeoMx Digital Spatial Profiler (DSP) profiles. (B) Relative cell viability of *EP300*-, *ARID1A*- and *CDH1*-perturbed cells treated with SAHA combined with XAV-939 (WNT pathway inhibitor) and palbociclib (CDK4/6 inhibitor) compared to SAHA alone in GES1 cells (n=3; mean ± SD) (*, p < 0.05, two-sided t-test). (C) Survival analysis of GCs with high and low expression of molecular phenotypes associated with SAHA-enriched clusters G1 and G4 in the ACRG and TCGA cohorts. Significance was determined using the univariate log-rank test and multivariate Cox proportional hazard model. (D) Stained GeoMx DSP slides annotated with low expression of SAHA-enriched clusters G1 (blue) and G4 (red), based on ssGSEA difference scores (G1-G4), for patient NGC512. Each circle within the stained slide represents a tumor ROI, annotated with ssGSEA difference scores.

Primary GCs are associated with high levels of inter- and intra-patient (epi)genomic heterogeneity, raising the possibility that some molecular phenotypes in our dcPerturb-seq compendium might be expressed in tumors even in the absence of overt epigenetic challenge. To investigate this possibility, we applied the dcPerturb-seq molecular signatures to primary GC expression profiles from the ACRG and TCGA cohorts (**Figure 4A**) [56, 57]. Primary GCs were stratified into high- and low-expressing groups based on the expression levels of the resistance-associated gene clusters. We found that decreased expression of the SAHA G1 and G4 gene clusters were consistently associated with significantly poorer survival in both the ACRG and TCGA cohorts, with survival associations remaining significant in multivariate analyses (p value < 0.05) (**Figure 4C**). Patients with high expression of GO pathways related to EMT and WNT signaling, observed in G1-perturbed cells *in vitro* (Figure 3C), also exhibited significant poorer survival (**Figure S8C**-**D)**. These results highlight the potential clinical utility of molecularly subtyping primary GCs using dcPerturb-seq molecular phenotypes.

As GCs are known to exhibit significant intra-patient heterogeneity [58], we also examined if dcPerturb-seq molecular phenotypes could be applied to improve our understanding of GC spatial heterogeneity. Applying the SAHA G1 and G4 gene clusters to an in-house spatial transcriptomic dataset using NanoString GeoMx Digital Spatial Profiler (GeoMx DSP) technology [14], we observed clear spatial distributions of tumor spatial regions of interest (ROIs) exhibiting distinct G1- and G4-low signatures within and across multiple patients (NGC512, Moran’s I = 0.24, p = 2.6 × 10^-4^; NGC510, Moran’s I = 0.48, p = 6.1 × 10^-4^; NGC520, Moran’s I = 0.30, p = 1.5 × 10^-6^; NGC531, Moran’s I = 0.50, p = 8.4 × 10^-4^) (see **Methods**) (**Figure 4D** and **Figure S8E**). Deconvolution analysis revealed comparable proportions of tumor and non-tumor cells within G1-low and G4-low ROIs (**Figure S8F**). This finding suggests that the expression of molecular phenotypes are not uniformly distributed across a tumor section.

### Cell-Extrinsic Therapeutic Pressures Reveal Additional Gene Perturbation/Molecular Phenotype Network Interactions

To explore how cell extrinsic stressors such as drug pressures might modulate gene-gene network interactions, we integrated the entire compendium of dcPerturb-seq molecular phenotypes across all five exposure conditions (DMSO control plus four epigenetic drugs), using a common analysis pipeline to quantify global transcriptomic changes between gene-perturbed and control cells under distinct epigenetic treatments (**Figure 5A**). Surveying ∼4.3 million interactions, we observed that transcriptomic shifts caused by the same gene perturbation were often altered, sometimes dramatically, in a drug-dependent manner. For example, in *KDM5A*-perturbed cells, we observed down-regulation of cell growth related processes following SAHA treatment, compared with down-regulation of apoptotic and cell-substrate adhesion pathways after 5-AZA treatment (**Figure S9**). To systematically examine how transcriptional responses to the same genetic perturbation can be modulated by epigenetic drug exposures, we quantified the number and identity of up-regulated genes either shared across multiple therapies or private to specific therapies (**Figure 5B**). This analysis revealed distinct patterns. Perturbation of *STK11*, a serine-threonine kinase commonly altered in various cancers [59], induced a broadly common set of changes across all five conditions, while *ESRRG* gene perturbation, a member of the estrogen receptor-related family, exhibited both distinct and shared transcriptional alterations across conditions. *PBRM1*, a remodeling complex encoding gene [60], showed strong drug-specific responses upon perturbation, possibly related to previously described context-specific functions of the SWI/SNF complex [61].

**Figure 5.**
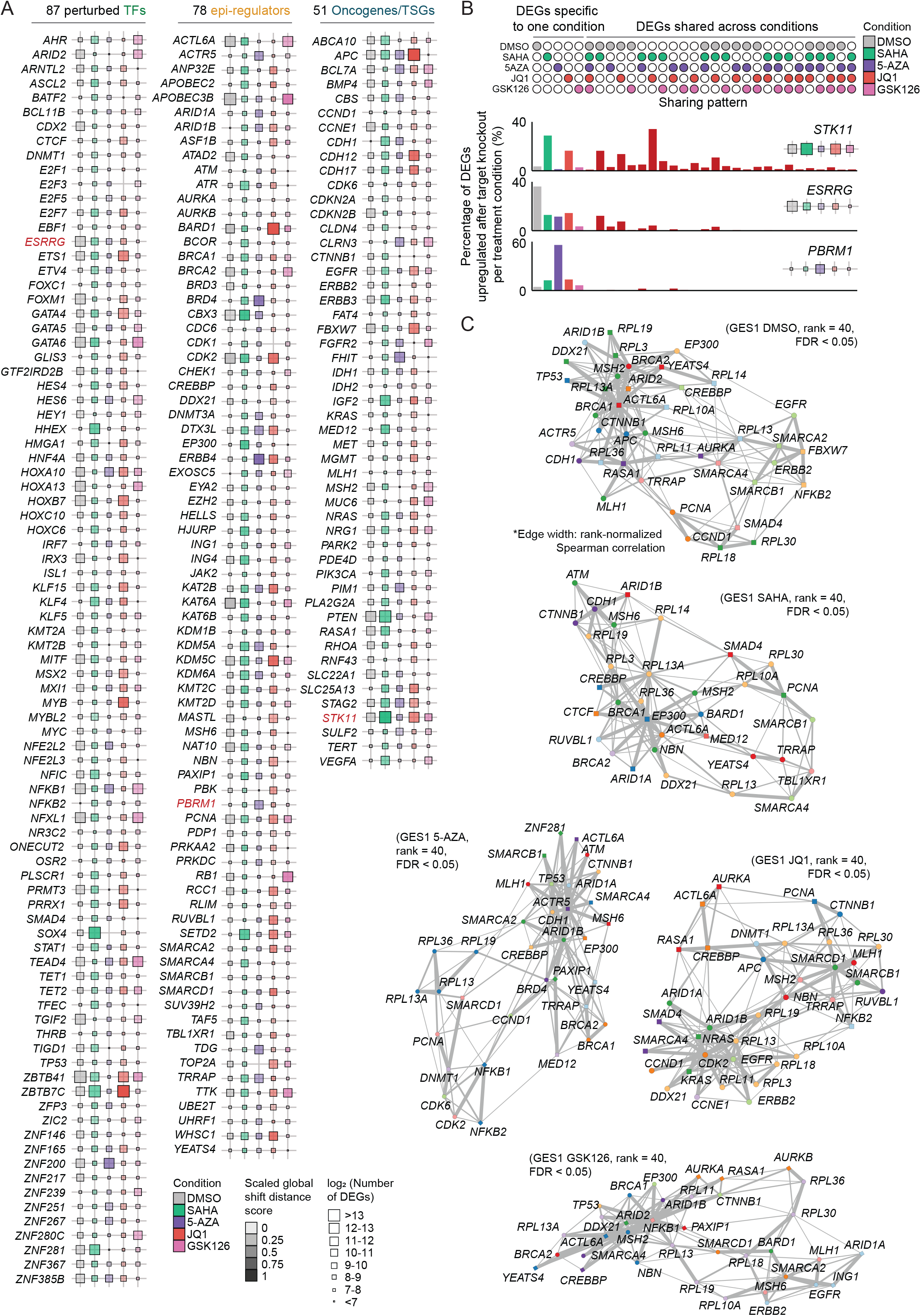
Epigenetic Drug Exposures Reveal Additional dcPerturb-seq Gene-Gene Network Interactions. (A) Graphic summary representing the magnitude of transcriptomic alterations for each dcPerturb-seq gene perturbation under DMSO and epigenetic drug exposure, relative to negative control cells. Each treatment was analyzed independently and represented by a distinct color. (B) Percentage of up-regulated DEGs (AD test, FDR < 0.05) following each drug treatment, grouped by sharing pattern. Genes upregulated in only one condition (left) or shared by two or more conditions (right). (Bottom) Examples of distributions associated with *STK11*, *ESRRG* and *PBRM1* perturbations. (C) Gene-gene interaction networks using correlations of mean molecular phenotype profiles from dcPerturb-seq under DMSO and epigenetic drug treatment. Edge width: rank-normalized Spearman correlation for dcPerturb-seq network (rank = 40, FDR < 0.05, permutation test with 10,000 random permutations).

To visualize the landscape of gene perturbation-epigenetic drug interactions from a network perspective, we integrated the dcPerturb-seq compendium with CORUM [39] and constructed gene-gene interaction networks [62]. Distinct networks were observed for each epigenetic treatment, indicating that drug exposures can uncover additional gene-gene interactions not overtly discernible in the absence of extrinsic therapeutic challenges (**Figure 5C**). For example, in DMSO-treated cells, chromatin remodeling genes (*ACTL6A* and *ARID2*) formed a regulatory core correlated with the DNA damage repair genes (*MSH2*, *MSH6*, *BRCA1* and *BRCA2*) (FDR < 0.05). However, under SAHA treatment, *EP300* emerged as a regulatory core together with *CREBBP*. Notably, since both *EP300* and *CREBBP* encode acetyltransferases [63], it is possible that perturbation of these two acetyltransferases may normalize abnormal SAHA-induced histone acetylation resulting in therapy resistance. In 5-AZA-treated cells, chromatin complex encoding genes (*ARID1A*, *ARID1B* and *ACTR5*) were enriched, consistent with **Figure 3E** where these three genes exhibited similar patterns of gene program and GO pathway regulation. Taken collectively, these results reveal how cell-extrinsic drug challenges can uncover novel gene-perturbation molecular phenotypes.

### Conserved and Context-Specific Perturbation Responses Under Basal and Extrinsic Drug Exposures Across Cell Types

Finally, to explore if dcPerturb-seq molecular phenotypes observed in GES1 cells are also present in other gastric cell lines, we performed dcPerturb-seq across another five lines. In two lines, we repeated all four epigenetic treatments SNU719 (192,778 cells) and NUGC3 (201,478 cells), and in three lines with SAHA only: LMSU (44,409 cells), SNU1750 (41,228 cells), and HGC27 (24,301 cells). Quality control of the additional dcPerturb-seq data was achieved similar to GES1 cells (**Figure S10A-B)**. The final data set encompasses 4,746 gene/drug/cell line multiplexed interactions across 625,866 cells in 6 gastric cell lines from diverse subtypes (**Supplementary Table 5**) covering 226 genes and 4 epigenetic drugs (**Figure S10C**).

To ask if genes likely to cause broad transcriptional impact in one line are also likely to cause broad impact in another, we focused on control (DMSO) conditions and computed shift distance scores across lines. Individual lines showed global positive correlations with median shift-distance scores across lines (**Figure 6A**), a finding corroborated by TI analysis (**Figure S10D**). Stratifying the targets by their variation in shift distance scores revealed that genes showing commonly broad transcriptional impact across lines (bottom 20% variance) exhibited higher transcriptional stability [64] and higher essentiality (measured by CRISPR screening) (**Figure 6B**). These low variance targets also exhibited lower variance in their regulation of downstream biological processes (**Figure 6C**), exemplified by genes such as *AURKA*, *CDK6* and *AURKB* in regulating survival-related pathways [65–67] (**Figure 6D**). In contrast, we also identified genes showing context- and cell-line specific effects (**Methods**) (dots highlighted in orange; **Figure 6A**; **Supplementary Table 6**). For instance, compared to other cell lines, *ZNF367* and *CTNNB1* perturbations caused strikingly high levels of transcriptional impact in SNU1750 and HGC27 cells respectively (**Figure 6E** and **Figure S10E**). Interestingly, SNU1750 and HGC27 cells also express the lowest levels of *ZNF367* and *CTNNB1* expression respectively, highlighting target gene basal expression as a determinant of transcriptional impact.

**Figure 6.**
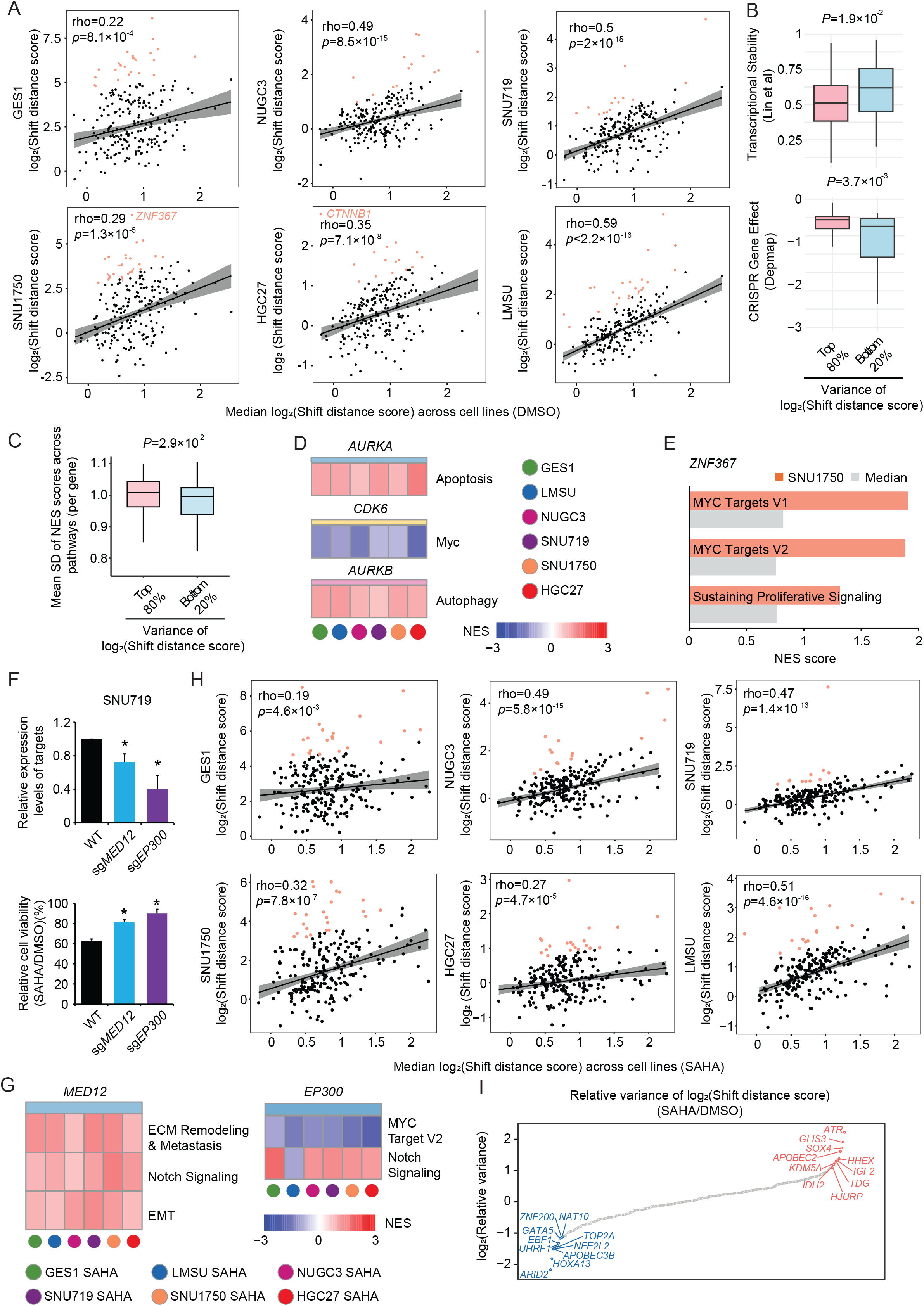
Conserved and Context-Specific Perturbation responses Across Cell Types Under Basal and Extrinsic Drug Exposures. (A) Shift distance scores for gene perturbations in individual cell lines versus median scores across six cell lines under DMSO control, with outliers (orange) annotated. Significance was determined using Pearson correlations. (B) Boxplots comparing transcriptional stability indexes (top; stably expression across species and developmental stages) and DepMap CRISPR gene effect scores (bottom; lower gene effect score indicated higher essentiality) for perturbations with bottom 20% and top 80% variance of log_2_(Shift distance score) across cell lines (*P* values are adjusted p values calculated by Wilcoxon rank sum test). (C) Boxplots comparing mean standard deviation (SD) of normalized enrichment scores (NES) across pathways for perturbations with bottom 20% and top 80% variance of log_2_(Shift distance score) across cell lines (*P* values are adjusted p values calculated by Wilcoxon rank sum test). (D) Heatmap of normalized enrichment scores (NES) for pathway analysis in *AURKA-*, *CDK6*- and *AURKB*-perturbed cells, compared to WT cells, across six gastric cell lines (GES1, LMSU, NUGC3, SNU719, SNU1750 and HGC27) under DMSO control. Normalized enrichment scores were generated from GSEA. (E) Normalized enrichment scores (NES) for pathway analysis in *ZNF367-*perturbed cells, compared to WT cells, for SNU1750 and the median across six gastric cell lines under DMSO control. Normalized enrichment scores were generated from GSEA. (F) qRT-PCR analysis of *MED12* and *EP300* gene expression levels in WT and *MED12-* and *EP300*-perturbed cells (top); Relative cell viabilities of WT, *MED12-* and *EP300-* perturbed cells treated with SAHA compared to DMSO in SNU719 cells (bottom) (n=3; mean ± SD) (*, p < 0.05, two-sided *t*-test). (G) Heatmap of normalized enrichment scores (NES) for pathway analysis in SAHA-treated *MED12-* and *EP300*-perturbed cells, compared to WT cells, across six gastric cell lines. Normalized enrichment scores were generated from GSEA. (H) Shift distance scores for gene perturbations in individual cell lines versus the median score across six cell lines under SAHA treatment, with outliers (orange) annotated. Significance was determined using Pearson correlations. (I) Distribution of relative variance of log_2_(Shift distance score) comparing SAHA-treated conditions to DMSO control for all genes targeted by the sgRNA library. Top (more context-specific) and bottom (more conserved) enriched perturbations are highlighted in orange dots and blue dots respectively.

To assess how extrinsic drug exposures might alter these conserved and context-specific molecular phenotypes, we analyzed SAHA-treated profiles across all six cell lines. Cell viability profiles remained significantly correlated (**Figure S11A**), mirroring conservations seen in DMSO conditions (**Figure S11B**). Specifically, *MED12*, *EP300,* and *KLF4* perturbations induced resistance to SAHA in multiple lines, a finding we further experimentally and independently validated in SNU719 cells (**Figure 6F** for *MED12* and *EP300,* similar to **Figure S5C**). Molecular phenotypes associated with certain gene perturbations were also conserved. For example, *MED12* perturbation in SAHA-treated cells resulted in upregulation of EMT processes across all six cell lines, while *EP300* perturbation led to the upregulation of Notch signaling (**Figure 6G**). Likewise, *KLF4* perturbation led to the upregulation of Myc targets, glucose metabolism and DNA repair pathways in multiple lines (**Figure S11C, Figure S11D** provides additional examples). Across the lines, shift distance scores per line largely correlated with median levels across cell lines under SAHA treatment, albeit with certain cell-specific outliers such as *SOX4* in GES1 cells (**Figure 6H**). We also assessed how SAHA challenge across different lines might impact variances in shift distance scores (**Figure 6I**). Interestingly, some perturbations such as *SOX4* and *APOBEC2* (which interact with HDAC transcriptional corepressor complexes [68, 69]), exhibited increased line-to-line variance upon SAHA treatment, indicating a shift towards context-specific dominance after HDAC inhibition. Conversely, other perturbations, such as *ARID2* and *APOBEC3B*, became more conserved under SAHA exposure. Taken collectively, these results reveal both conserved and context-specific gene-perturbation molecular phenotypes associated with cell-extrinsic drug challenges.

## Discussion

In this study, we applied dcPerturb-seq to interrogate >200 GC-associated genes, systematically characterizing their downstream molecular phenotypes in the context of epigenetic drug pressure. Clinical management of GC remains highly challenging, with current therapeutic strategies often limited by the emergence of drug resistance [70]. While some targeted therapies have demonstrated efficacy [71, 72], GC treatment options remain limited. GCs also exhibit substantial genomic heterogeneity that can contribute to metastasis and terminal resistance. Notably, compared to the many putative GC targets identified by genomic sequencing and transcriptomic profiling studies [57, 73, 74], comparatively little is known about the functional pathways regulated by these targets. Here, after providing confidence in the reliability of our dcPerturb-seq platform by multiple lines of stringent QC, we observed high concordance with previously characterized gene-specific molecular phenotypes, such as *FBXW7* in mTOR signaling regulation [75], and *KDM5C* in WNT signaling [76]. Further, by interpreting the dcPerturb-seq molecular phenotypes as a network, we gained a deeper understanding into the potential cellular functions of poorly characterized genes such as *ZBTB41*. Previous studies have posited a relationship between *ZBTB41* expression and the p53 signaling, and *ZBTB41* has been proposed as biomarker for cervical cancer [77]. These findings provide initial inroads into the subsequent study of frequently altered GC genes, and this data set which is publicly available should thus provide a valuable resource for future research.

Functional cancer phenotypes are often influenced by GxG and GxD interactions. For example, HER2-positive GCs with co-occurring alterations in cell cycle components (eg *CCNE1*) have impaired responses to trastuzumab [78], and in colorectal cancer *KRAS* mutations can act as drivers of acquired resistance to anti-EGFR therapy [79]. Novel molecular phenotypes have also been shown to arise from the cooperative activity on multiple oncogenic alterations, beyond a purely additive model [19]. Technically, the need to explore multiple GxG and GxD combinations poses significant challenges for traditional bulk RNA sequencing (RNA-seq), which require each perturbation to be performed as a separate experiment. Advancements in Perturb-seq technologies offer unprecedented potential to dissect complex GxG and GxD interactions at the single-cell level. Previous Perturb-seq studies have investigated transcriptional networks governing tissue development, immune responses, and cancer progression [23, 80, 81], however these have typically focused on a limited number of genes (<50), primarily to validate gene sets identified through genome-wide bulk CRISPR screens. More recently, genome-scale Perturb-seq screens have demonstrated the capacity to assign gene functions and dissection of complex cellular phenotypes [7, 82], however these studies have largely focused on methodological development. To our knowledge, the current study is one of the first to study therapy resistance in cancer and the first to apply Perturb-seq to GC.

We investigated if single-cell CRISPR molecular phenotyping can be used to dissect gene dosage-dependent effects, in a manner not achievable using bulk RNA-seq. Notably, we observed several dosage-sensitive (DS) genes among our targets, including well-characterized haploinsufficient genes such as *KAT6A*, *APC* and *GATA6*. Specifically, APC haploinsufficiency has been linked to impaired hematopoiesis [83], while GATA6 haploinsufficiency is associated with pancreatic agenesis [84]. Our findings also revealed a separate population of genes exhibiting a dosage-linear (DL) pattern, aligning with previous findings for genes such as *JAK1* and *IRF1*, which demonstrate gradual changes in transcriptional regulation with increasing degrees of perturbation [85]. While our dataset did not reveal a distinct dosage-tolerant (DT) pattern, we speculate that this may be attributed to limitations in cell number or our selection of target genes. Overall, these observations emphasize the highly nuanced relationship between gene dosage and transcriptional regulation, which single-cell CRISPR screening is uniquely positioned to explore.

By integrating the dcPerturb-seq platform with drug exposures, we identified targets exhibiting resistance and sensitivity to epigenetic treatment. Reassuringly, our dcPerturb-seq molecular phenotypes recapitulated several published findings, such as *MED12* loss inducing an EMT-like molecular signature, leading to chemotherapy resistance in colon cancer and resistance to EGFR-inhibitor in lung cancer [86]. We found that SAHA (HDACi) resistance could be caused by *EP300* perturbation. In breast cancer, *EP300* loss has been associated with paclitaxel resistance [54]. As another finding, *KLF4* deficiency in our study was also associated with HDACi resistance, and *KLF4* has been linked to increased tamoxifen resistance in breast cancer [87]. Besides targets where perturbation leads to resistance, our study also revealed targets conferring sensitivity upon perturbation, such as *FOXM1* and *KMT2B*, suppression of which sensitized cancer cells to chemotherapy [88, 89]. These may also represent promising avenues for further investigation in the context of synthetic lethality.

Our ability to link molecular phenotypes with functional cell viability yielded insights into heterogeneous pathways associated with epigenetic drug resistance. We found that diverse molecular pathways can converge to produce a common physiological phenotype (drug resistance). This is conceptually akin to melanoma cells exhibiting mesenchymal-like, dedifferentiated, and metabolically reprogrammed “starved-like” states after MAPK inhibition, demonstrating the coexistence of multiple drug-tolerance transcriptional circuits [90]. In our data, the SAHA_G1 resistance gene cluster exhibited enriched molecular phenotypes associated with phenotypic plasticity, such as epithelial-to-mesenchymal transition (EMT), WNT signalling and tissue development processes. Such plasticity, including epithelial-to-mesenchymal transition (EMT), lineage transdifferentiation, and reversion to stem or progenitor-like states [27], have been shown to strongly correlate with transcriptional plasticity and treatment resistance [16, 91–93]. In contrast to SAHA_G1, the SAHA_G4 resistance gene cluster showed enrichment of molecular signatures associated with cell cycle, metabolic, and stress-related responses, all of which have also been linked to therapy resistance. For example, heterogeneity in cell cycle behaviour has been identified as a critical determinant to chemotherapy resistance [94], and enrichment of carbohydrate and nucleotide metabolism pathways have been observed in PARP-inhibitor-resistant ovarian cancer cells [95].

The data richness of our dcPerturb-seq molecular phenotyping dataset allowed us to examine the complex interplay between epigenetic drug treatment and genetic perturbations. Specifically, our analysis revealed that certain gene perturbations, such as *STK11*, *PCNA*, *TEAD4*, and *ZNF165*, tended to elicit broadly similar responses across treatment conditions, while other perturbations (e.g., *PBRM1*, *TP53*, *SMARCA4*, and *SMAD4*) elicited more treatment specific molecular phenotypes. Epigenetic challenge also enabled the emergence of distinct gene-gene regulatory networks not observable in the absence of drug exposure. For example, JQ1 treatment highlighted two dominant regulatory cores: genes encoding components of the SWI/SNF complex (*SMARCD1* and *SMARCB1*) and the *CDK2* gene, while GSK126 challenge uncovered an enriched core of genes (*NFKB1*, *MSH2*, *ARID2*, *ACTL6A* and *ARID1B*), associated with downregulation of stress-related pathways, reminiscent of phenotypic inertia [81]. Both conserved and context-specific patterns were observed in our comparative analysis across cell lines in terms of genetic perturbations. It is important to note, however, that such cell line specificity likely arises from a complex interplay of genomic, epigenetic, transcriptomic, and metabolic factors, underscoring the challenge of fully dissecting the mechanistic basis of these context-specific responses to genetic perturbation, particularly following therapeutic exposure.

This study has limitations and can be expanded in many directions. Incorporating additional time points will enable the tracking of longitudinal changes in molecular phenotypes, as drug resistance is a gradual and multifaceted process [96]. Extending the duration of drug treatment may facilitate the detection of induced mutational signatures, as observed in solid tumors receiving epigenetic therapy [97]. The accuracy and generalizability of our findings can be further enhanced by incorporating additional cell lines, increasing the number of replicates, cells per condition, and sequencing depth. A more comprehensive understanding of the role of chromatin modifiers will inevitably require investigation at the chromatin level. Techniques such as SHARE-seq [98] or other single-cell multi-omic approaches will be crucial for elucidating the intricate interplay between chromatin structure and cellular function. Finally, experimental costs can be further reduced by exploring alternative scRNA-seq methods, such as scifi-RNA-seq [99] and SPLiT-seq [100]. Looking ahead, the development of *in vivo* spatial single-cell screening techniques [101] will be crucial for recapitulating the complex cellular stromal/immune interactions and environmental challenges encountered during tumor growth which are absent in conventional cell line models.

In conclusion, when benchmarked against key single-cell perturbation datasets catalogued by scPerturb.org [102] (**Figure S11E**), our dcPerturb-seq dataset is comparably larger in scale and unique in its combination of genetic and pharmacological perturbations, enabling the study of extensive combinatorial interactions. Information provided through such platforms may accelerate the development of foundational and large-language AI models for cancer, as exemplified by recent single-cell foundation models such as scGPT [103], GET [104], and virtual cell model STATE [105], to predict cellular responses to perturbations and infer gene regulatory networks. These findings will contribute significantly to data-driven resources for translational cancer research, paving the way for the development of more effective and personalized therapeutic strategies.

## Methods

### Targets selection for CRISPR library construction

To prioritize targets for the dcPerturb-seq assay, we curated a candidate gene list based on their potential to significantly influence transcriptional regulation. We began with a compiled list of 1,639 human transcription factors (TFs) [106] and 720 epigenetic regulators [107]. Separately, we compiled 105 oncogenes, tumor suppressor genes (TSGs), and epigenetic regulators frequently altered in cancer, as supported by existing literature [56, 57, 108]. After removing duplicates, this initial set contained 2,290 targets. We then determined which of these targets were expressed in gastric cell lines (our experimental model). Using bulk RNA-seq data from 90 gastric cell lines, we filtered for targets with detectable expression (log2(TPM+1) > 0), narrowing the list to 1,915 genes. This refined list was then subjected to differential expression analysis in primary gastric cancer datasets (TCGA and local cohort Singapore cohort #1-SG1). Applying a threshold of absolute fold change > 1.5 and an adjusted p-value < 0.01 in both cohorts identified 120 significantly dysregulated targets. The 105 literature-curated cancer genes were exempt from this differential expression filter, as their relevance in cancer progression is well-established despite the potential lack of significant transcriptomic perturbation.The final candidate list of 216 targets was then confirmed to be detectable in scRNA-seq data to ensure compatibility with our dcPerturb-seq output.

### Cell culture, dcPerturb-seq CRISPR library design and lentiviral production

SNU719 and SNU1750 cells were obtained from Korean Cell Line Bank. NUGC3 cells were obtained from the Japan Health Science Research Resource Bank. HGC27 cells were obtained from CLS Cell Lines Service. LMSU cells were obtained from Riken Cell Bank. Cell line identities were confirmed by STR DNA profiling at the Centre for Translational Research and Diagnostics, National University of Singapore. All lines were negative for mycoplasma contamination as assessed by the MycoAlert™ Mycoplasma Detection Kit (Lonza). Culture medium information and latest mycoplasma test dates are summarized in **Supplementary Table 5.**

CRISPR sgRNA libraries were synthesized by Sigma using scaffolds (U6-gRNA-10X:EF1a-puro-2A-Cas9-2A-tGFP) containing Capture Sequences compatible with 10x Genomics Feature barcoding technology (**Supplementary Table 1**). Single guide RNAs containing oligos were cloned into lentiCRISPR v2 backbones for independent validation. Lentivirus particles were produced by co-transfecting HEK293T cells with transfer plasmids and standard packaging vectors. For dcPerturb-seq, cells were infected with pools of lentiviral particles containing four independent sgRNAs per target gene, and also non-targeting sgRNAs as controls. Cells were transduced with lentiviral pools at an MOI of 0.2 to attain single sgRNA entries per cell, and selected with puromycin to achieve a final library representation of roughly 2,000 cells per sgRNA. Lentivirus infected GES1, SNU719 and NUGC3 cells were recovered after 2 weeks,and subsequently treated with SAHA (SML0061, Merck), 5-AZA (A3656, Merck), JQ1 (11187, Cayman Chemical) and GSK126 (S7061, Selleck Chemicals) for an additional 2 weeks before proceeding to single cell profiling. Similarly, SNU1750, HGC27 and LMSU cells were treated with SAHA (SML0061, Merck) for 2 weeks before undergoing single cell profiling. Briefly, dcPerturb-library infected cells were seeded at approximately 80% confluency and cultured in medium containing the compound at the specified concentration. The drug-containing medium was refreshed every 48-72 hours. Cells were monitored and passaged upon reaching confluency. This cycle of treatment and passaging was repeated for the entire two-week duration. Drug concentrations for all cell lines are presented in **Supplementary Table 7**.

### dcPerturb-seq library preparation and sequencing

Cells were prepared for single-cell RNA-sequencing in 1×PBS with 0.1% BSA as detailed in the 10x Genomics Single Cell Protocols Cell Preparation Guide (10x Genomics, CG000053 Rev C). Cells were separated into droplet emulsions using the Chromium Controller (10x Genomics) and the 10x Genomics Chromium Single Cell 3[ Reagent Kits v3.1 with Feature Barcode Library Kit (10x Genomics, PN-1000121 and PN-1000079), with the goal of recovering 10,000 cells per GEM group before filtering. For preparation of gene expression and sgRNA libraries, samples were processed according to 10x Genomics Chromium Next GEM Single Cell 3[ Reagent Kits v3.1 User Guide with Feature Barcode technology for CRISPR Screening (CG000205 Rev D). Sequencing libraries were generated using unique sample indexes, which were quantified using the Kapa library kit. mRNA and sgRNA libraries were pooled at a 4:1 ratio and sequenced on both Illumina Hiseq4000 and NovaSeq X plus sequencing platforms (Illumina) according to the 10x Genomics User Guide.

### Cell cycle analysis

Cell cycle distribution was analyzed by flow cytometric quantification of DNA content using propidium iodide (PI) staining. Briefly, adherent cells were harvested via trypsinization, and washed in ice-cold PBS. Cell pellets were resuspended in PBS and fixed by the addition of ice-cold 100% ethanol to a final concentration of ∼70% by vortexing. Fixed cells were stored overnight at -20°C. Subsequently, cells were washed twice with PBS and treated with RNase A (R1253, Thermo Fisher Scientific; 0.1 mg/mL in PBS) for 1 hour at 37°C to degrade RNA. Cells were stained with PI (P1304MP, Thermo Fisher Scientific; 100 µg/mL) overnight at 4°C. Data acquisition was performed on LSR Fortessa (BD Biosciences) and analyzed using FlowJo (v10.10.0).

### Cell viability and Quantitative Real-Time PCR assays

Cell viability was assessed using the Cell Counting Kit-8 (CCK-8; Dojindo) according to the manufacturer’s instructions. Briefly, cells were seeded in 96-well plates and treated with the indicated concentrations of epigenetic drugs for 72 hours. For combinatorial treatments, cells were incubated for 72 hours with either SAHA (SML0061, Merck; 1.25μM) alone or in combination with one of the following compounds: XAV-939 (HY-15147, MedChemExpress; 1μM), palbociclib (HY-50767, MedChemExpress; 2μM), DAPT(HY-13027, MedChemExpress; 2μM), semagacestat (HY-10009, MedChemExpress; 4μM), vactosertib (HY-19928, MedChemExpress; 8μM), or galunisertib (HY-13226, MedChemExpress; 4μM). Viability was measured following this incubation period. For quantitative Real-Time PCR assays, cells were seeded in 6-well plates and treated with epigenetic drugs for 72 hours before proceeding to sample collection for downstream analysis. Total RNA was extracted using RNA extraction reagents (Qiagen), and reverse transcribed using iScript Reverse Transcriptase Supermix (Biorad). Quantitative real-time PCR was performed using SYBR Green PCR Master Mix (Life Technologies). Samples were analyzed using the Bio-Rad CFX96 system. Experimental Ct values were normalized to *GAPDH*, and relative mRNA expression was calculated as fold changes compared to *GAPDH* expression (2-ΔCt). Sequences of primers are listed in **Supplementary Table 8**.

### dcPerturb-seq data processing

Cell Ranger 6.1.2 software (10x Genomics) was used for alignment of scRNA-seq reads to human transcriptomes, alignment of sgRNA reads to the dcPerturb-seq library, collapsing reads to UMI counts, and cell calling. The 10x Genomics GRCh38 version 2020-A genome build was used as a reference transcriptome. Subsequently, Seurat V4.3.0 [109] was utilized to perform basic quality control (QC) filtering. To filter out low-quality cells and to retain the most informative genes, we filtered out cells with >12% (HFE145), >20% (GES1), >25% (SNU719), >25% (NUGC3), >12% (SNU1750), >12% (HGC27) and >12% (LMSU) mitochondrial UMIs per cell. We then removed cells with RNA counts below the 1st percentile or above the 99th percentile and genes with counts below the 1st percentile. The threshold number of guides was set to 2-10 based on the threshold plot, and cells with only one guide were kept. Each cell was categorized according to its guide identity representing a single genetic perturbation. Only cells bearing a single guide were used for downstream analysis. The data was then normalized using Seurat (Relative Counts normalization, RC-norm). After RC-normalization, genes were further filtered by being highly expressed (> 2 after RC-norm) in no less than five cells. The data was then analysed for highly variable genes (HCG) using the FindVariableFeatures() function, scaling across cell populations using the ScaleData() function, and dimensional reduction analysis using PCA.

To evaluate the overall expression profiles of each perturbed target gene and to remove possible background effects, we first calculated the mean expression of all cells on the whole transcriptome for each perturbed target gene. We then applied Z-transformation on the average gene expression matrix using mean expression and standard deviation values for negative control sgRNA cells as the standard reference. Correlations between any two perturbed target genes was calculated based on these Z-transformed vectors.

### Single cell long-read RNA-seq library preparation, sequencing and data processing

Libraries for single-cell long-read RNA-seq were prepared from amplified cDNA (10x Genomics Chromium Next GEM Single Cell 3’ Kit v3.1) using the Kinnex single-cell RNA kit and sequenced using PacBio long-read technology by Next Level Genomics. HiFi reads were generated and analyzed by the SMRT analysis pipeline (https://www.pacb.com/products-and-services/analytical-software/smrt-analysis/). Specifically, HIFI reads were de-concatemerized with skera. Adaptor/primer removal and demultiplexing were performed with lima. Full-length non-concatemer reads were processed with isoseq refine and clustered with isoseq cluster2. Clustered FLNC reads were aligned to GRCh38 using pbmm2, producing BAM result files. For downstream per-read indel parsing we retained primary, mapped alignments and excluded secondary/supplementary records. “Deletion reads” were defined as reads containing ≥1 CIGAR D event overlapping a pre-specified CRISPR edited interval (BED) for the target gene; CIGAR N (introns) did not count as deletions. Cell barcodes were extracted from PacBio reads from the CB tags and matched to a 10x cell barcodes whitelist. Classified deletion reads were linked to target genes via this barcode–guide map. To ensure accuracy, targets with <10 total reads overlapping the edited interval were excluded. For each target, the deletion rate was (# reads with D overlapping the edited interval)/(# reads overlapping the interval), and the fold change (FC) was computed as FC = deletion_rate_target / deletion_rate_negative_control. All analyses of BAMs used pysam 0.23.3; BED intersections used bedtools 2.30.0.

### CRISPR-mediated gene alteration impact

To evaluate the impact of CRISPR-mediated gene alterations, we compared the expression of each target gene in cells with corresponding sgRNA to the cells with non-targeting control guides. To determine if CRISPR guides targeting the same gene produced similar transcriptional impacts on target gene, we compared the expression of the intended target gene in guide-perturbed cells versus all other cells for each target gene. Differential expression was computed with Seurat’s FindMarkers() function, which applies the Wilcoxon rank-sum test to unpaired groups. To ensure robustness, we repeated the analysis after applying additional filters: the bottom 25% expressed targets and the targets with bottom 25% cell numbers were excluded. The results remained consistent, confirming the reliability of our findings (**Figures S12A-D**).

### Shift distance test and differential gene expression analysis using AD test and TRADE

To assess the impact of gene perturbations on molecular phenotypes, we employed energy-based statistical tests (Hotelling’s *T*^2^ test) as Oana et al [31] to calculate “shift distance scores”. These tests are well-suited for comparing distributions in high-dimensional data, making them ideal for analyzing gene expression profiles represented as vectors of principal component scores. Specifically, each cell was characterized by its top 20 principal components, and we evaluated whether the distribution of these vectors differed between control cells and those subjected to gene perturbations. Divergences in these distributions indicate that the perturbation alters the structure or distribution of transcriptional profiles within perturbed cells.

Recognizing the inherent biological variability of gene expression, the incomplete penetrance of certain perturbations, and the heterogeneity of some gene expression programs, we adopted a non-parametric approach. This approach circumvents the need for specific distributional assumptions about gene expression. Specifically, gene expression was z-normalized relative to control cells. For each gene, we tested whether the distribution of normalized expression values was identical between control cells harboring non-targeting sgRNAs and cells subjected to each gene perturbation. The Anderson-Darling test [7] was utilized for this purpose due to its broad sensitivity to differences in distributions, allowing detection of subtle changes in expression profiles.

To assess the global transcriptional impact of each perturbation, we also performed differential gene expression analysis using TRADE, quantifying effects via the transcriptome-wide impact (TI) score as defined by Ajay et al [32]. Briefly, differential expression was performed on a pseudobulk matrix in which sequencing lanes were first aggregated, and TI was calculated with TRADE (version 0.99, default settings). TI scores were log-transformed to stabilize variance. To validate the robustness of our results, we repeated the TI-related correlation analysis while excluding the lowest-expressed 25% of targets and targets present in the lowest 25% of cells and observed consistent findings (**Figures S12E**). To determine whether a high TI reflected strong effects on a few genes or weaker effects on many, we examined the distribution of “valid DEG numbers” named Π_DEG_ reported by TRADE. Using established thresholds (Π_DEG_ = 45 for all genes, 500 for essential genes; Ajay et al.), we found 74/226 perturbations (32.7%) showed broad effects (≥ 45 Π_DEG_) while 91/226 (40.3%) exhibited focused effects. To examine if sgRNAs targeting the same gene showing correlation in TI scores, we performed one-way ANOVA tests after filtering the sgRNAs with the bottom 25% target expression and lowest 25% of cell numbers. Finally, to assess perturbation impact of targets stratified by LOEUF scores, we calculated the ratio of the mean TI of targets within the curated LOEUF groups to the overall mean TI across all target perturbations.

To identify genetic perturbations with context-specific impact, we compared their shift distance score/TI in each cell line to the median score across all six lines. A linear model was fit to this relationship, and perturbations residing above the upper bound of the 80% prediction interval were classified as context-specific.

### Minimum-Distortion Embedding (MDE) based 2D-Projection

Minimum-distortion embedding (MDE) has been used in single-cell data to identify connections of different elements using high-dimensional gene profiling data [7]. Using the Z-transformed gene profiling vectors of each target gene, we first used PyMDE (https://github.com/cvxgrp/pymde) to generate a 20 dimension spectral embedding (affinity=’nearest_neighbors’, n_neighbors=7, eigen_solver=’arpack’), and then performed dimension reduction to 2D for visualization (n_neighbors=7, constraint=pymde.Standardized(), repulsive_fraction=5). PyMDE was run multiple times in parallel to find the optimal distortion and residual norm values. The embeddings were plotted as graphs using the pymde.plot() function, and then manually annotated using Gene Ontology and CORUM database annotation.

### Permutation test for *RPL* genes

To evaluate the significance of the essentiality of *RPL* genes for cell survival and proliferation, we randomly selected 40 negative control sgRNAs, matching the number of sgRNAs targeting *RPL* genes. We then calculated the number of cells containing the selected negative control sgRNAs and the sgRNAs targeting *RPL* genes. To determine the significance of enrichment, we performed 10,000 random samplings. An empirical P-value was derived by counting the number of instances in which the number of cells containing the randomly selected negative control sgRNAs was smaller than the number of cells containing sgRNAs targeting *RPL* genes.

### CRISPR viability scores

For each condition (DMSO, SAHA, JQ1, 5-AZA, and GSK126), we determined the number of cells assigned to a specific sgRNA-targeted gene. This count was normalized by dividing it by the total number of cells in the respective condition and multiplying the result by 10^5^. To mitigate computational errors, a pseudo-cell count was added to the value. The resulting metric represents the number of cells containing a specific sgRNA-targeted gene per 10^5^ cells. To calculate CRISPR viability scores for a specific gene perturbation under a drug condition, we computed the log_2_ fold change of the resulting metric between the drug and DMSO conditions. Finally, CRISPR viability scores were z-normalized by subtracting the mean and dividing by the standard deviation of the scores across all sgRNA-targeted genes. When calculating CRISPR viability scores using sgRNA counts, we replaced the cell numbers with the counts of sgRNAs targeting a specific gene. This substitution allowed us to use sgRNA numbers as the basis for determining CRISPR viability scores.

### Gene dosage effect and cell cycle analysis

For each target gene, we used the RC-normalized gene expression profiles and defined cells with zero expression of the gene as “complete gene removal” (CR), and defined cells that still have intermediate expression of the gene as “partial gene removal” (PR). For each target gene, we merged the cells with negative control cells, then reperformed the normalization step on the merged cells, using Seurat to perform log-normalized raw counts, found variable features (n = 1000), and data scaling to enhance variability. Differential gene expression analysis was performed between negative control cells and fully depleted cells using the Seurat FindAllMarkers() function (min.pct = 0.25, logfc.threshold = 0.25). Identified DEGs with adjusted p-value <=0.05 were retained. The expression patterns of negative control cells, partially depleted cells and fully depleted cells were visualized using the R package Pheatmap.

We used two methods to model gene dosage effect, namely the Kullback–Leibler divergence (KL divergence) based method, and the Euclidian distance-based method. For these two methods, we calculated the distances between the “negative control - PR” and the “CR - PR” separately for each target gene using the expression profiles (log-normalized) of DEGs (between negative control and CR) for all cells, and then calculated differences between the distances, forming a distribution of the distances for classifying the dosage effect subtypes.

#### For the KL Divergence based method

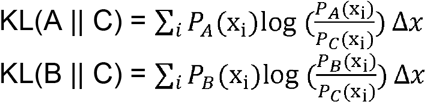

Differences = KL(A || C) - KL(B || C)

Where *P_A_(x) , P_B_(x), P_C_ (x)* denotes estimated probability through Kernel Density Estimation (KDE) on gene expression values of negative control cells (A), completely depleted target gene KO cells (B) and partially deplete target gene KO cells (C). x_i_ denotes the i-th point on a common grid of values (determined from the KDE).

#### For the Euclidean Distance based method

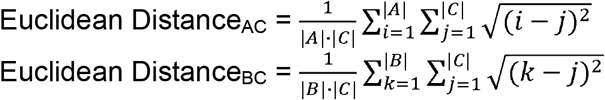

Differences = Euclidean Distance_AC_ - Euclidean Distance_BC_

Where A represents gene expression values of negative control cells, B represents gene expression values of completely depleted target gene KO cells (CR), and C represents gene expression values of intermediate-depleted target gene KO cells (PR). ∣A∣, ∣B∣ and ∣C∣ denote the sizes of sets A, B, and C, respectively. I, j and k represent a gene expression value in a single cell in A, B and C, respectively.

#### Determination of dosage effect categories

Using differences in either KL Divergence or Euclidean Distance for multiple DEGs, we first estimated the probability through Kernel Density Estimation (KDE) on all DEGs as P(x). Then we determined the category (DS, DL, or DT) of the dosage effect of a target gene by checking the Area Under Curve (AUC) patterns.

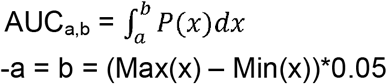

-a = b = (Max(x) – Min(x))*0.05

Where a, b were determined by the differences between maximum and minimum values.

Dosage-Sensitive (DS): AUC_b,max(x)_ = Max(AUC_min(x),a_, AUC_a,b_ , AUC_b,max(x)_) Dosage-Linear (DL): AUC_a,b_ = Max(AUC_min(x),a_, AUC_a,b_ , AUC_b,max(x)_) Dosage Tolerant (DT): AUC_min(x),a_ = Max(AUC_min(x),a_, AUC_a,b_ , AUC_b,max(x)_)

Cell circle analysis was performed using the Seurat function CellCycleScoring, which used default s.genes and g2m.genes from references [110]. Cell cycle proportion changes were visualized using the R package ggplot2.

### Clustering of gene expression programs and GO biological pathways

For the top resistance-associated target genes (n = 50), we first excluded target genes which have less than 35 cells. Z-transformed gene expression profiles for the remaining target genes were then used for unsupervised clustering using the R package pheatmap (distance method is ‘Euclidean’, clustering method is ‘ward.D2’). The number of clusters was manually decided based on their profiles on the heatmap.

We performed pathway enrichment analysis using Gene Set Enrichment Analysis (GSEA) with the GO Biological Process (GO BP) database. To generate the ranked gene list, we calculated a sum statistic defined as sign (mean expression change) ×−log^10^ (P-value). The mean expression change for a gene was calculated as the difference between the mean expression of cells containing the sgRNA for the target gene and the mean expression of cells containing Negative Control sgRNAs. P-values were determined using the Anderson-Darling test. Cluster-enriched pathways were identified by comparing the normalized enrichment scores (NES) of pathways within the selected gene cluster to the NES of pathways in the other gene clusters. Pathways with a false discovery rate (FDR) of <0.001 and a mean difference >0.1 in NES scores were considered significant.

## Survival analysis

300 GC patients in the ACRG cohort and 415 GC patients in the TCGA cohort were collected to study associations between expression levels of dcPerturb-seq resistance molecular phenotypes and patient outcome. The expression levels of selected gene signatures were calculated using ssGSEA. We selected the top 25% of patients with the highest ssGSEA score of the gene signature as the “ Expression_high” group, and the bottom 25% of patients with the lowest ssGSEA score of the gene signatures as the “Expression_low” group. We employed Kaplan–Meier survival analysis with the overall survival as the outcome metric. Log-rank tests were used to estimate the significance of Kaplan–Meier curves. We established multivariate Cox proportional-hazards models using the R package “survminer.” Gene expression of the ACRG cohort was profiled using Affymetrix Human Genome U133 plus 2.0 Array (GSE62254, https://www.ncbi.nlm.nih.gov/geo/query/acc.cgi?acc=GSE62254) [56]. RNA-seq profiles of 415 primary GCs from the TCGA cohort were obtained from the Broad Institute TCGA Genome Data Analysis Center (GDAC) Firehose (https://gdac.broadinstitute.org/).

## Gene-Gene Interaction Network Analysis

Using gene-gene interactions of all possible pairs of perturbed target genes based on Z-transformed gene expression profile vectors, we employed a published software (https://github.com/robinmeyers/pan-meyers-et-al) [62] to construct gene-gene interaction networks based on the CORUM database (https://mips.helmholtz-muenchen.de/corum/). Target genes that are not in the CORUM database were excluded from analysis. Default software parameters were used. We selected rank 40 to visualize the network.

## Functional similarity analysis

To apply resistance associated molecular phenotypes from SAHA to other drug conditions/ cell lines/ cohorts, we applied single-sample gene set enrichment analysis (ssGSEA) algorithm using the R package GSVA. Gene names of the target genes in each group were used directly as ssGSEA input signatures. To identify enriched GO pathway signatures for each group, we performed ssGSEA for the GO pathways and then calculated average scores according to the identified GO pathway signatures.

## DSP analysis

Collection and bioinformatics pre-processing on the DSP GC dataset was performed according to previous studies [14, 111]. Cell type proportion of the DSP data was estimated by CIBERSORTx using paired scRNA-seq samples. Similar to methods of applying the identified resistance signatures from SAHA to other drug conditions/ cell lines/ cohorts, ssGSEA was used to map signatures to DSP tumor regions of interest (ROI). Spatial autocorrelation of the difference in ssGSEA scores between the G1 and G4 clusters within each primary GC sample was evaluated using Moran’s I test, as implemented in the R package spdep (v1.2-8).

## Data availability

The raw data generated for this study has been deposited in NCBI SRA XXXXX.

## Contributors

Conceptualisation: CX, HM, TS and PT; data acquisition: CX, MHL, SABAG, SSL, SS, XO, STT, SWTH, ALKT, FZ, MT, RS, WPY, JBYS; formal analysis: CX, HM, TS, LPJ, KKH, OZX, DRP; Facilities, reagents and intellectual support: MHL, XO, STT, ALKT, HA, ASLC, ADJ, SL, JR; drafting of manuscript: CX, HM, TS and PT.

## Grant support and Acknowledgements

This research was supported the National Medical Research Council grant MOH-000967. This work was also supported by the National Research Foundation, Singapore, and Singapore Ministry of Health’s National Medical Research Council under its Open Fund-Large Collaborative Grant (“OF-LCG”) (MOH-OFLCG18May-0003). This research was also supported by the Duke-NUS Core funding. We thank Duke-NUS Genome Biology Facility (DGBF) for supporting reagent procurement and preparation of sequencing libraries.

