## Supplementary Figures for "Diverse Pharmacogenomic Resistance Landscapes Across Epigenetic Drug Classes Revealed by Gastric Cancer Perturb-seq"

Chang Xu^1*^, Haoran Ma^1*^, Taotao Sheng^2*^, Ming Hui Lee^1^, Siti Aishah Binte Abdul Ghani^1^, Laura Perlaza-Jimenez^3,4^, Kie Kyon Huang^1^, Shen Kiat Lim^1^, Supriya Srivastava^5^, Xuewen Ong^1^, Su Ting Tay^1^, Shamaine Wei Ting Ho^2^, Ong Zhi Xuan^1^, Angie Lay Keng Tan^1^, Feng Zhu^5^, Hassan Ashktorab^6^, Alfred Sze-Lok Cheng^7^, Anand D. Jeyasekharan^5,8,9,10^, Shang Li^1,11^, Ming Teh^12^, Raghav Sundar^1,13,14^, David R. Powell^4^, Joseph Rosenbluh^3,15^, Wei Peng Yong^8,9^, Jimmy Bok-Yan So^10,13,14,16,17^, Patrick Tan^1,2,8,11,14,18,19,#^

**Affiliations**

^1^Cancer and Stem Cell Biology Program, Duke-NUS Medical School, Singapore

^2^Genome Institute of Singapore, Agency for Science, Technology and Research, Singapore

^3^Cancer Research Program and Department of Biochemistry and Molecular Biology, Biomedicine Discovery Institute, Monash University, Clayton, VIC, Australia

^4^Bioinformatics Platform, Monash University, Clayton, VIC, Australia

^5^Department of Medicine, Yong Loo Lin School of Medicine, National University of Singapore, Singapore

^6^Department of Medicine, Howard University, Washington, DC, USA

^7^School of Biomedical Sciences, The Chinese University of Hong Kong, Hong Kong, China

^8^Cancer Science Institute of Singapore, National University of Singapore, Singapore

^9^Department of Haematology-Oncology, National University Health System, Singapore, Singapore

^10^NUS Centre for Cancer Research, Yong Loo Lin School of Medicine, National University of Singapore, Singapore, Singapore

^11^Department of Physiology, Yong Loo Lin School of Medicine, National University of Singapore, Singapore

^12^Department of Pathology, Yong Loo Lin School of Medicine, National University of Singapore, Singapore

^13^Yong Loo Lin School of Medicine, National University of Singapore, Singapore

^14^Singapore Gastric Cancer Consortium, Singapore

^15^Functional Genomics Platform, Monash University, Clayton, VIC, 3800, Australia

^16^Department of Surgery, University Surgical Cluster, National University Health System, Singapore

^17^Division of Surgical Oncology, National University Cancer Institute, Singapore

^18^SingHealth/Duke-NUS Institute of Precision Medicine, National Heart Centre Singapore, Singapore

^19^Cellular and Molecular Research, National Cancer Centre, Singapore

*Equal Contribution

**^#^Corresponding author**

Prof Patrick Tan

Cancer and Stem Cell Biology Program, Duke-NUS Medical School, Singapore

**Supplementary Figures**
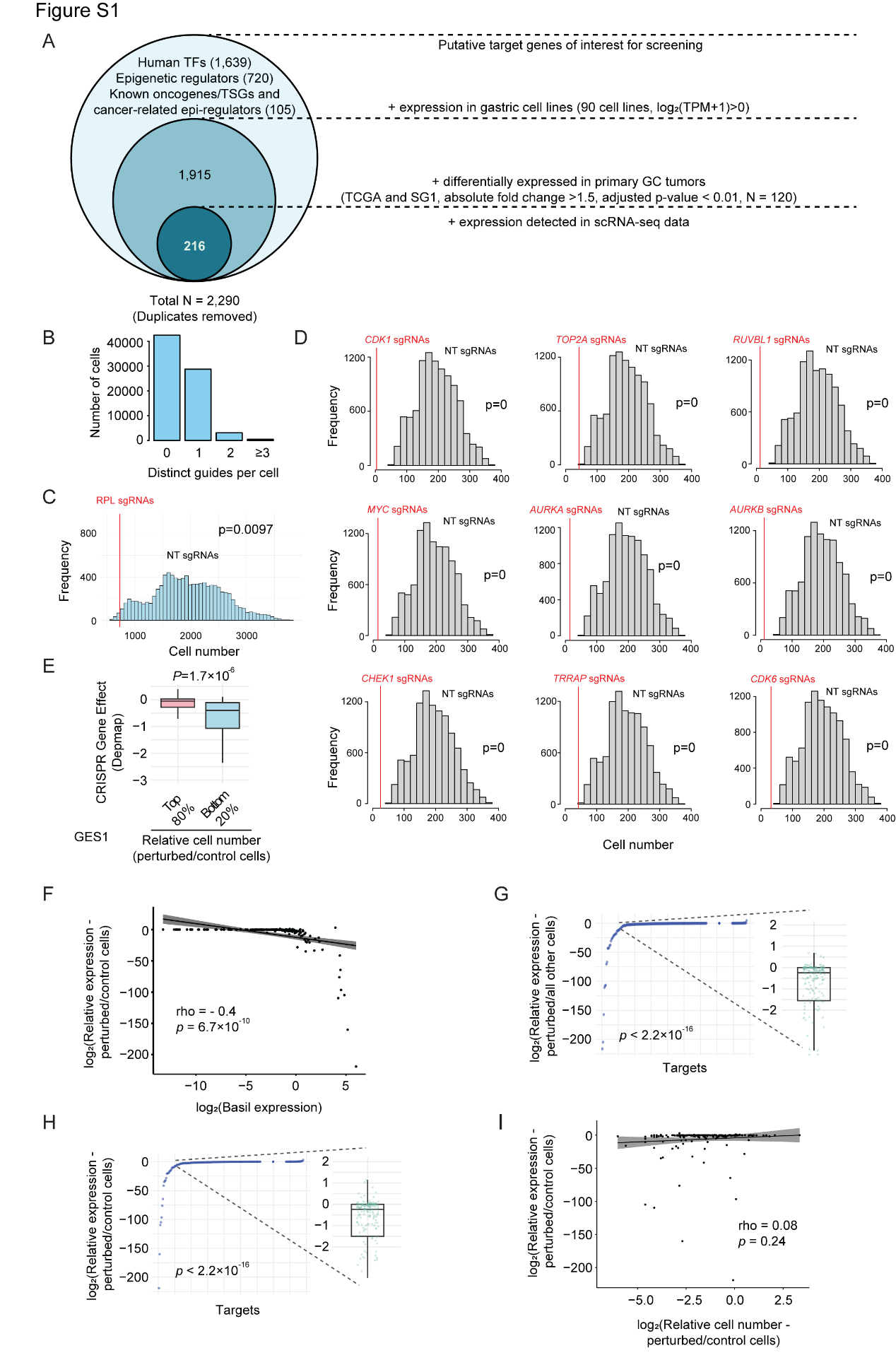

**Supplementary Figure1 (Related to Figure 1)**

(A) Schematic representation of the target gene selection process for dcPerturb-seq library construction.

(B) Distribution of number of cells with 0, 1, 2, and ≥3 distinct guides captured per cell in GES1 cells.

(C) Comparison of cell number distributions between sgRNAs targeting *RPL* genes and randomly selected sets of non-targeting sgRNAs. Significance was determined by an empirical permutation test.

(D) Comparison of cell number distributions between sgRNAs targeting common essential genes predicted by Depmap and randomly selected sets of non-targeting sgRNAs. Significance was determined by an empirical permutation test.

(E) Boxplots comparing DepMap CRISPR gene effects for perturbations with the bottom 20% and top 80% relative cell number (perturbed/control cells) (*P* values are adjusted p values calculated by Wilcoxon rank sum test).

(F) Correlation between basal expression levels and relative target expression (perturbed/control cells) for each target gene in GES1 cells. Significance was determined using Pearson correlations.

(G) Distribution of log_2_(relative target expression) for each target gene comparing perturbed cells with all other cells. Significance was determined using binomial tests.

(H) Distribution of log_2_(relative target expression) for each target gene comparing perturbed cells with control cells. Significance was determined using binomial tests.

(I) Correlation between relative cell number (perturbed/control cells) for each target gene and relative target expression (perturbed/control cells) in GES1 cells. Significance was determined using Pearson correlations.

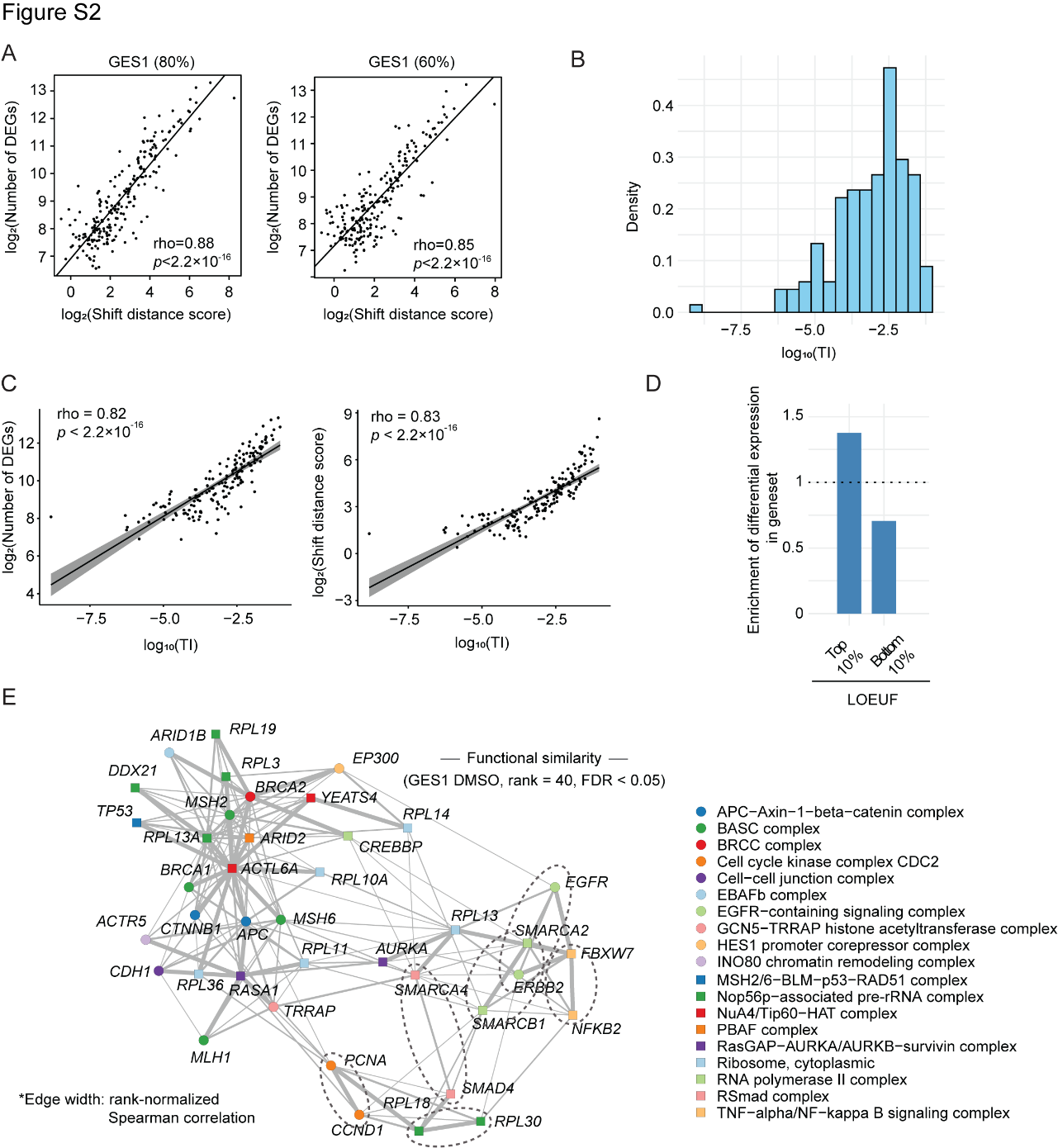

**Supplementary Figure 2 (Related to Figure 1)**

(A) Downsampling analysis examining correlations between shift distance scores relative to wild-type (WT) cells and number of differentially expressed genes (DEGs) for each target gene in 80% and 60% of down-sampled GES1 cells. Significance was determined using Pearson correlations.

(B) Distribution of log_10_(TI) for each target gene in GES1 cells.

(C) Correlation between TI and numbers of differentially expressed genes (DEGs) (left); Correlation between TI for each target gene and shift distance scores relative to wild-type (WT) cells (right) in GES1 cells. Significance was determined using Pearson correlations.

(D) Barplot plot illustrating enrichment of TI scores in target genes stratified by loss-of-function intolerance (LOEUF). Genes with higher evolutionary constraint (LOEUF) exhibited greater enrichment of TI.

(E) Gene-gene interaction networks of perturbations in GES1 cell line, based on correlations in mean dcPerturb-seq expression profiles. Edge width: rank-normalized Spearman correlation for dcPerturb-seq network (rank = 40, FDR < 0.05, permutation test with 10,000 random permutations).

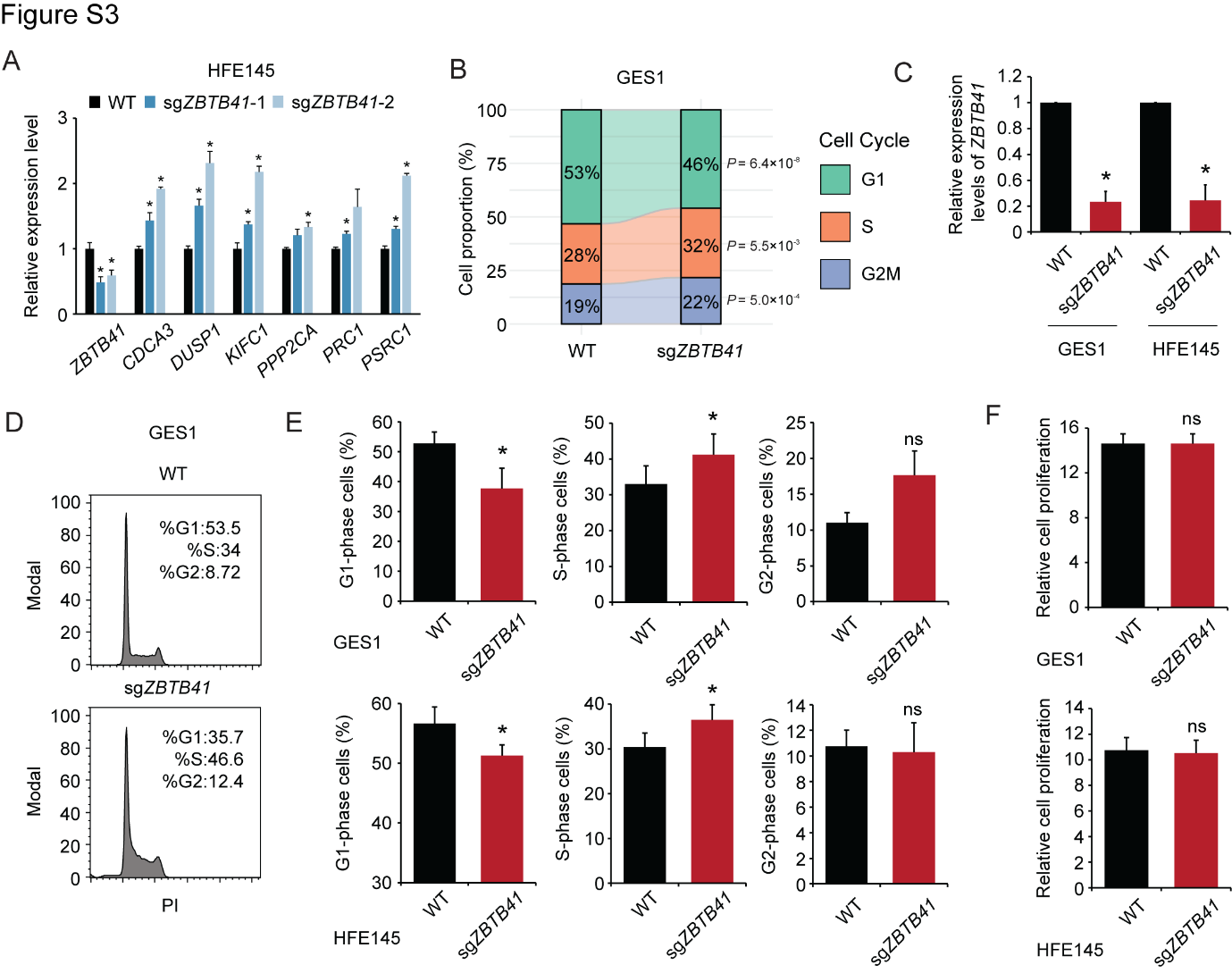

**Supplementary Figure 3 (Related to Figure 1)**

(A) qRT-PCR results showing relative expression levels of *ZBTB41*, *CDCA3*, *DUSP1*, *KIFC1*, *PPP2CA*, *PRC1* and *PSRC1* genes in WT and *ZBTB41*-perturbed HFE145 cells (n=3; mean ± SD) (*, p < 0.05, two-sided *t*-test).

(B) Comparison of cell cycle phase proportions (G1, S, G2M) in WT and *ZBTB41*-perturbed GES1 cells using dcPerturb-seq data. Significance was determined using Wilcoxon rank sum tests.

(C) qRT-PCR results of relative *ZBTB41* gene expression levels in WT and *ZBTB41*-perturbed GES1 and HFE145 cells (n=3; mean ± SD) (*, p < 0.05, two-sided *t*-test).

(D) Representative flow cytometry plots from cell cycle analysis of WT and *ZBTB41*-perturbed GES1 cells.

(E) Cell cycle phase proportions (G1, S, G2) in WT and *ZBTB41*-perturbed GES1 and HFE145 cells, as determined by flow cytometry (*, p < 0.05, two-sided *t*-test).

(F) Relative cell proliferation of WT and *ZBTB41-*perturbed GES1 and HFE145 cells (n=3; mean ± SD) (*, p < 0.05, two-sided *t*-test).

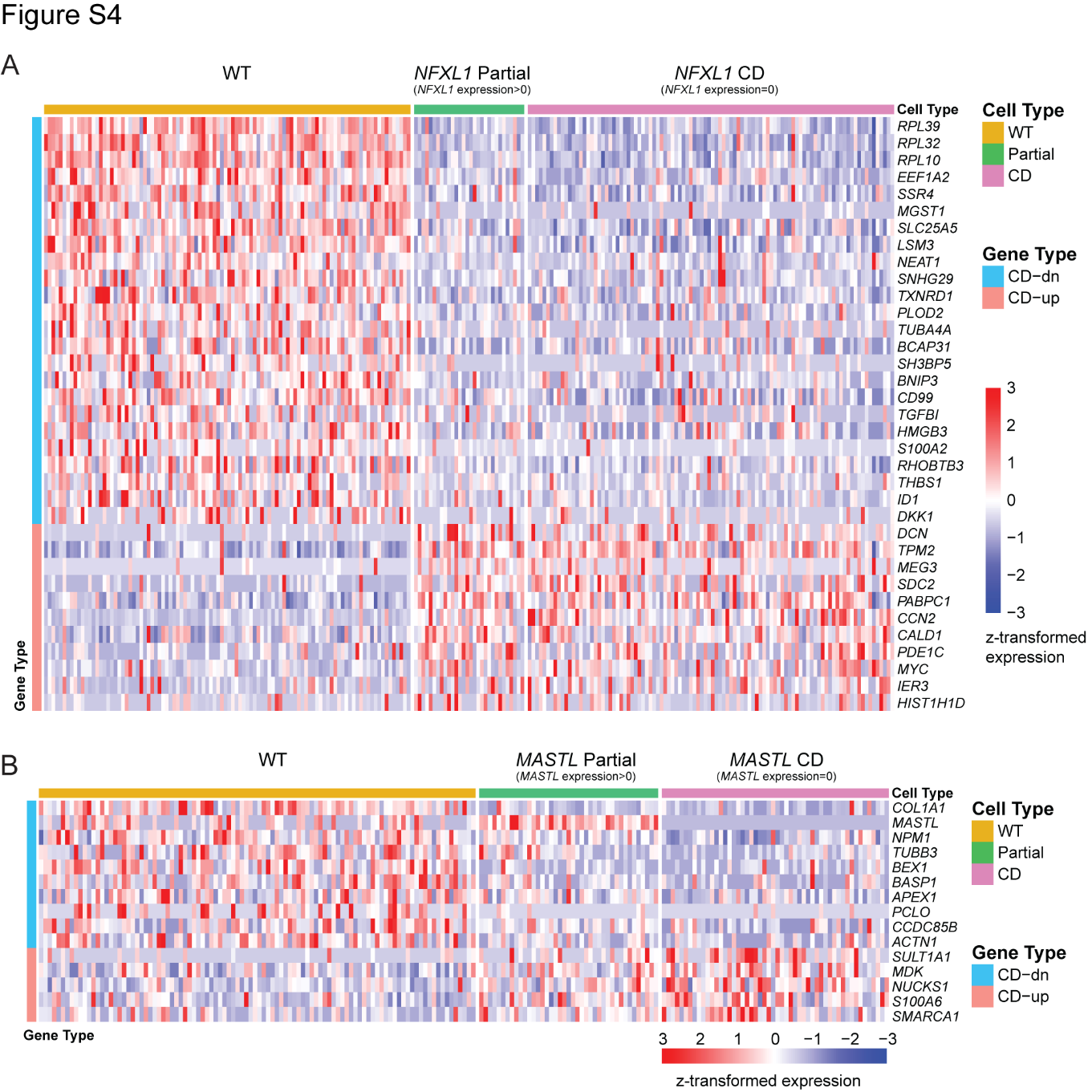

**Supplementary Figure 4 (Related to Figure 2)**

(A) Heatmap of downstream gene expression in WT, *NFXL1* partially-depleted (Partial) (expression) and completely-depleted (CD) cells. Columns represent cells belonging to either WT, *NFXL1* Partial or *NFXL1* CD groups. Rows represent genes differentially expressed between *NFXL1* CD and WT cells, identified by Wilcoxon rank sum tests. Row colors denote the direction of regulation in the *NFXL1* CD group relative to WT cells (blue, downregulated; orange, upregulated).

(B) Heatmap of downstream target gene expression in WT, *MASTL* partially-depleted (Partial) (expression) and completely-depleted (CD) cells. Columns represent cells belonging to either WT, *MASTL* Partial or *MASTL* CD groups. Rows represent genes differentially expressed between *MASTL* CD and WT cells, identified by Wilcoxon rank sum tests. Row colors denote the direction of regulation in the *MASTL* CD group relative to WT cells (blue, downregulated; orange, upregulated).

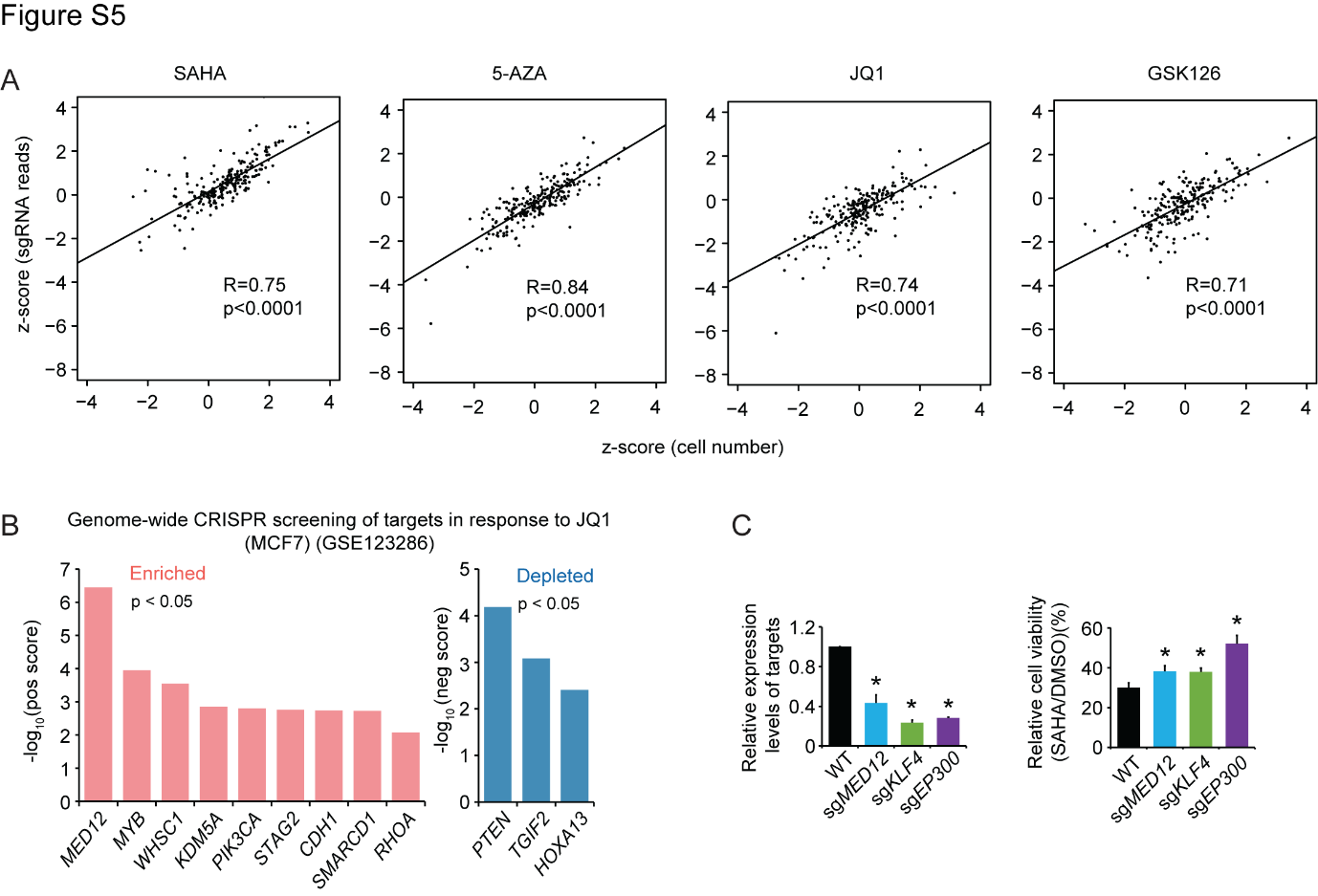

**Supplementary Figure 5 (Related to Figure 3)**

(A) Correlation between z-transformed cell numbers and z-transformed sgRNA read counts. Significance was determined using Pearson correlations.

(B) Representative target genes exhibiting significant enrichment or depletion in response to JQ1 treatment in MCF7 cells identified based on negative or positive effect scores using MAGeCK (p < 0.05). MCF7 data is from GSE123286.

(C) qRT-PCR analysis of relative expression levels of *MED12*, *KLF4*, and *EP300* in WT and *MED12*, *KLF4*, and *EP300* perturbed cells (left); Relative cell viability of WT and *MED12*, *KLF4*, and *EP300* perturbed cells treated with SAHA (1.25μM) compared to DMSO is shown (right) (GES1) (n=3; mean ± SD) (*, p < 0.05, two sided *t*-test).

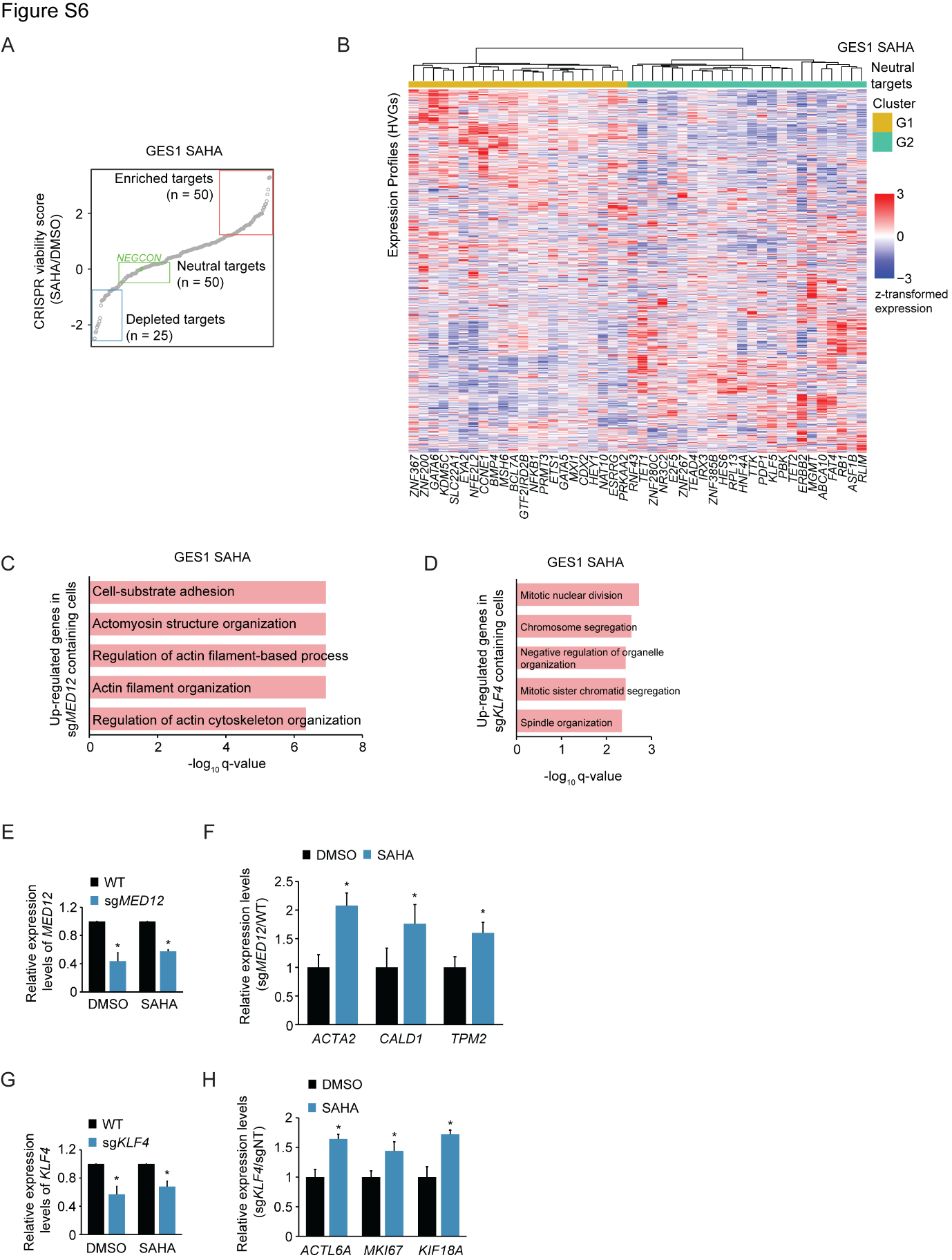

**Supplementary Figure 6 (Related to Figure 3)**

(A) Diagram illustrates the classification of gene perturbation targets as enriched, neutral, or depleted based on normalized CRISPR viability scores.

(B) Hierarchical clustering of neutral targets based on z-transformed expression profiles in response to SAHA treatment. Rows represent individual SAHA-neutral target gene perturbations; columns represent highly variable genes (HVGs) defined across the SAHA-neutral targets, at the level of dcPerturb-seq molecular phenotypes.

(C) Biological pathway enrichment analysis of genes perturbed by sgRNAs targeting *MED12* in GES1 cells under SAHA treatment. Values on the x-axis represent log_10_(q-value), where q-values are BH-adjusted p-values from the pathway enrichment tests.

(D) Biological pathway enrichment analysis of genes perturbed by sgRNAs targeting *KLF4* in GES1 cells under SAHA treatment. Values on the x-axis represent log_10_(q-value), where q-value are BH-adjusted p-values from pathway enrichment tests.

(E) qRT-PCR results of relative *MED12* gene expression levels in WT and *MED12*-perturbed GES1 cells under DMSO and SAHA treatment (n=3; mean ± SD) (*, p < 0.05, two-sided *t*-test).

(F) qRT-PCR analysis of relative expression levels of *ACTA2*, *CALD1*, and *TPM2* genes in *MED12*-perturbed cells compared to WT cells under DMSO and SAHA treatment (GES1) (n=3; mean ± SD) (*, p < 0.05, two-sided *t*-test).

(G) qRT-PCR results of relative *KLF4* gene expression levels in WT and *KLF4*-perturbed GES1 cells under DMSO and SAHA treatment (n=3; mean ± SD) (*, p < 0.05, two-sided *t*-test).

(H) qRT-PCR analysis of relative *ACTL6A*, *MKI67*, and *KIF18A* gene expression levels in *KLF4*-perturbed GES1 cells compared to WT cells under DMSO and SAHA treatment (n=3; mean ± SD) (*, p < 0.05, two-sided *t*-test).

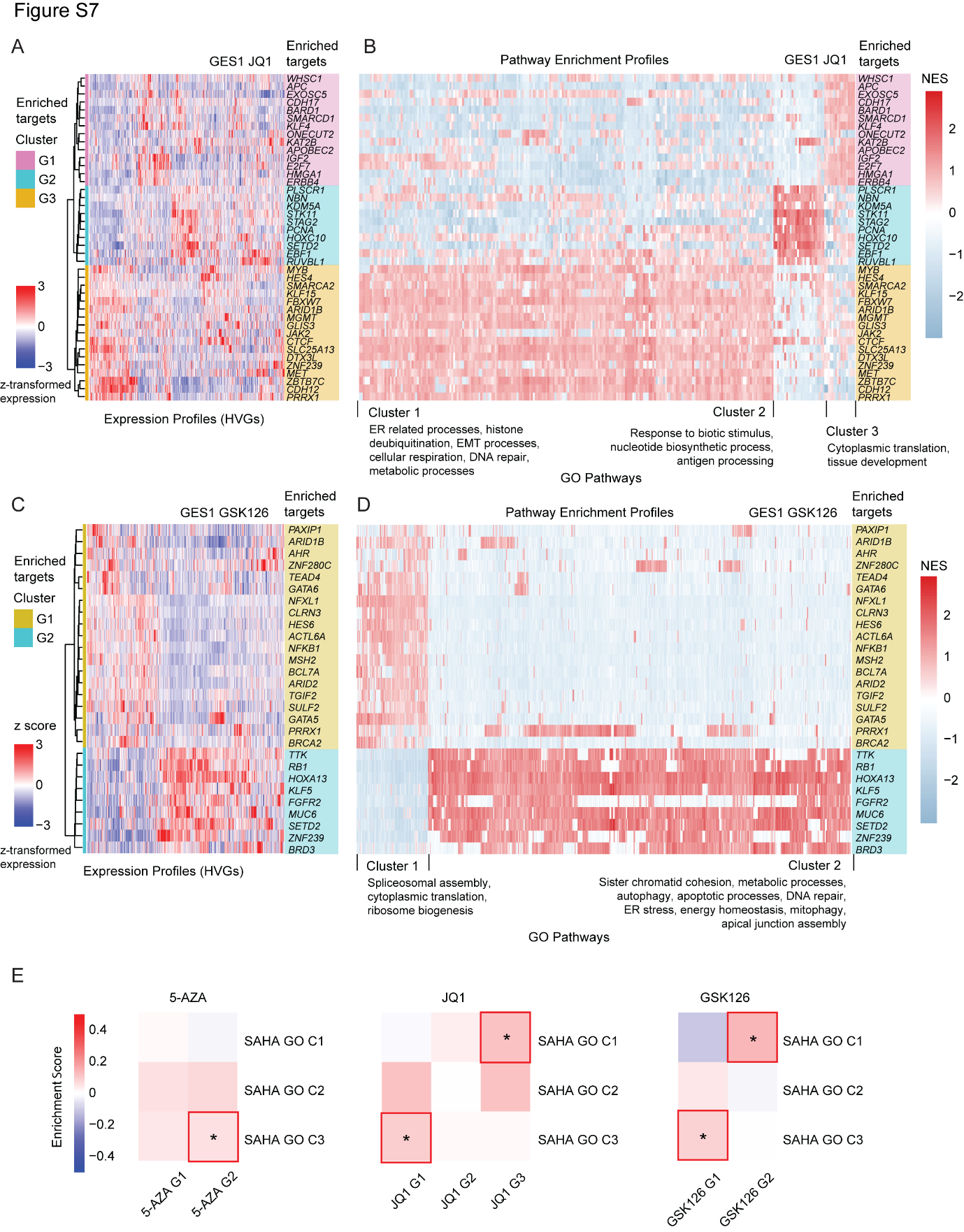

**Supplementary Figure 7 (Related to Figure 3)**

(A) Hierarchical clustering of enriched targets in response to JQ1 treatment based on z-transformed expression profiles. Rows represent individual JQ1-resistant target gene perturbations; columns represent highly variable genes (HVGs) defined across the JQ1-enriched targets, at the level of dcPerturb-seq molecular phenotypes.

(B)Heatmap illustrating NES scores for GO pathways associated with JQ1-enriched targets. Rows represent individual JQ1-enriched target gene perturbations in the same order and cluster as in **Figure S7A**; Columns represent GO pathways. Pathways were clustered by NES values and annotated by their cellular functions.

(C) Hierarchical clustering of enriched targets in response to GSK126 treatment based on z-transformed expression profiles. Rows represent individual GSK126-enriched target gene perturbations; columns represent highly variable genes (HVGs) defined across the GSK126-enriched targets, at the level of dcPerturb-seq molecular phenotypes.

(D) Heatmap illustrating NES scores for GO pathways associated with GSK126-enriched targets. Rows represented individual GSK126-enriched target gene perturbations in the same order and cluster as in Figure S7C; Columns represented GO pathways. Pathways were clustered by NES values and annotated by their cellular functions.

(E) Enrichment analysis of pathways enriched upon SAHA treatment across transcriptional profiles following 5-AZA, JQ1 and GSK126 treatments (*, p value < 0.05 by Wilcoxon rank sum test).

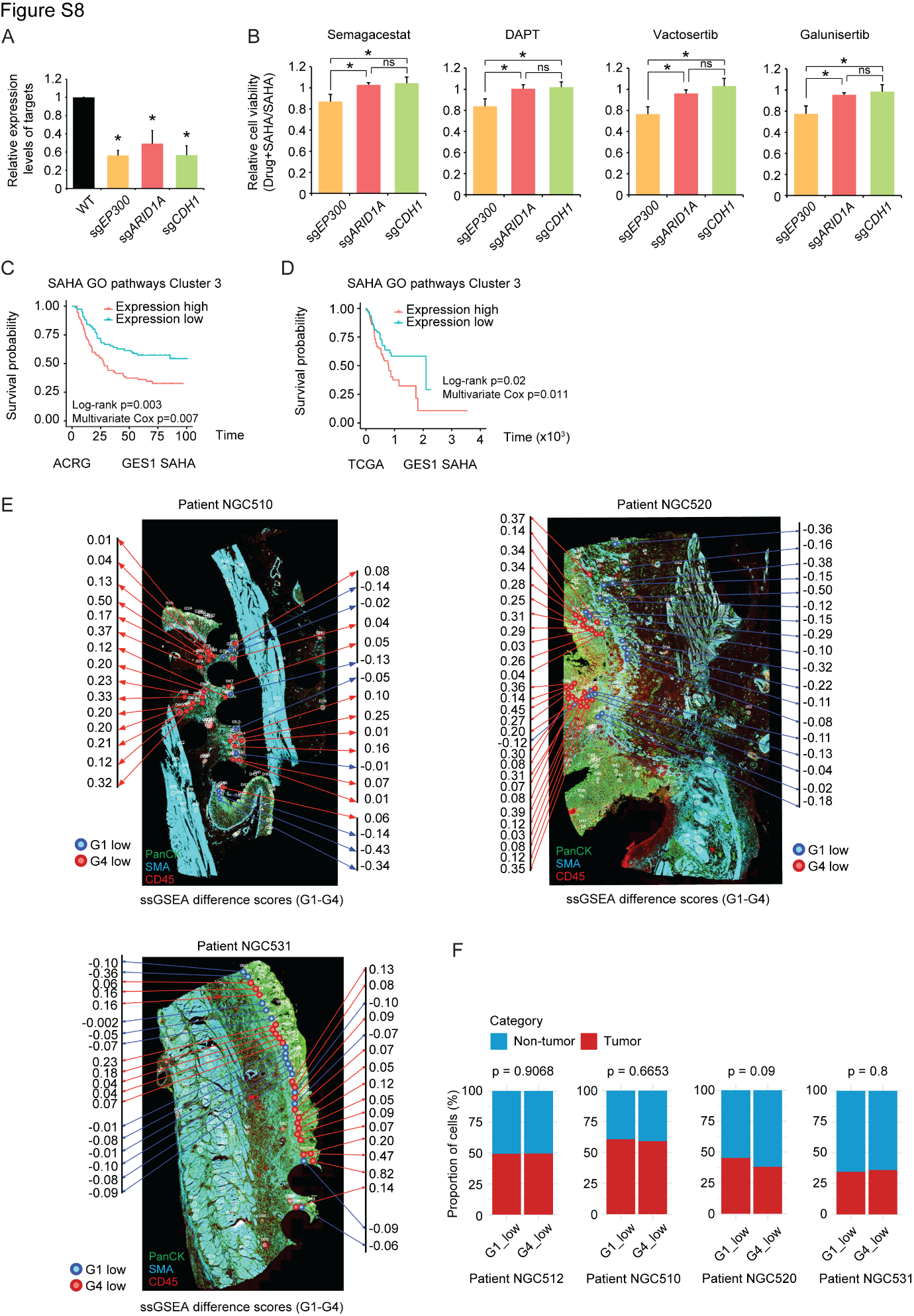

**Supplementary Figure 8 (Related to Figure 4)**

(A) qRT-PCR analysis of relative expression levels of *EP300*, *ARID1A*, and *CDH1* in WT and *EP300*, *ARID1A*, and *CDH1* perturbed cells (GES1) (n=3; mean ± SD) (*, p < 0.05, two-sided *t*-test).

(B) Relative cell viability of *EP300*-, *ARID1A*- and *CDH1*- perturbed cells treated with SAHA combined with semagacestat (Notch signaling inhibition), DAPT (Notch signaling inhibition), vactosertib (TGFβ inhibitor) and galunisertib (TGFβ inhibitor) compared to SAHA alone in GES1 cells (n=3; mean ± SD) (*, p < 0.05, two-sided t-test).

(C) Survival analysis of GCs with high and low expression of SAHA-enriched GO pathway cluster 3 in the ACRG cohort. Significance was determined using the univariate log-rank test and multivariate Cox proportional hazard model.

(D) Survival analysis of GCs with high and low expression of SAHA-enriched GO pathway cluster 3 in the TCGA cohort. Significance was determined using the univariate log-rank test and multivariate Cox proportional hazard model.

(E) Stained GeoMx DSP slides annotated with low expression of SAHA-enriched clusters G1 (blue) and G4 (red), based on ssGSEA difference scores (G1-G4), for patient NGC510, NGC520 and NGC531. Each circle within the stained slide represents a tumor ROI, annotated with ssGSEA difference scores.

(F) Cell proportion of non-tumor and tumor cells within ROIs in patient NGC512, NGC510, NGC520 and NGC531. Significance was determined using Wilcoxon rank sum test.

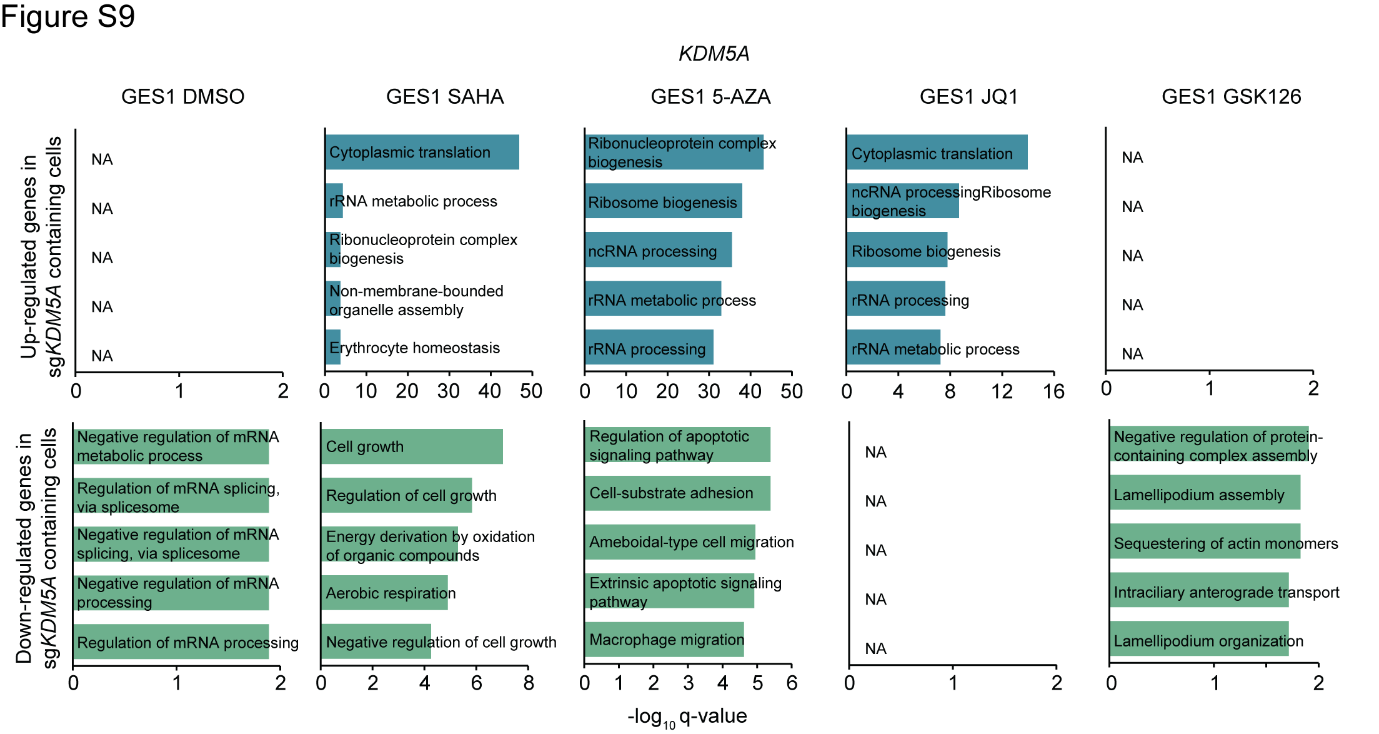

**Supplementary Figure 9 (Related to Figure 5)**

Biological pathway enrichment analysis of genes perturbed by sgRNAs targeting *KDM5A* in the GES1 cell line under DMSO, SAHA, 5-AZA, JQ1 and GSK126 treatment. Values on the x-axis represent log_10_(q-value), where q-values are BH-adjusted p-values from pathway enrichment tests.

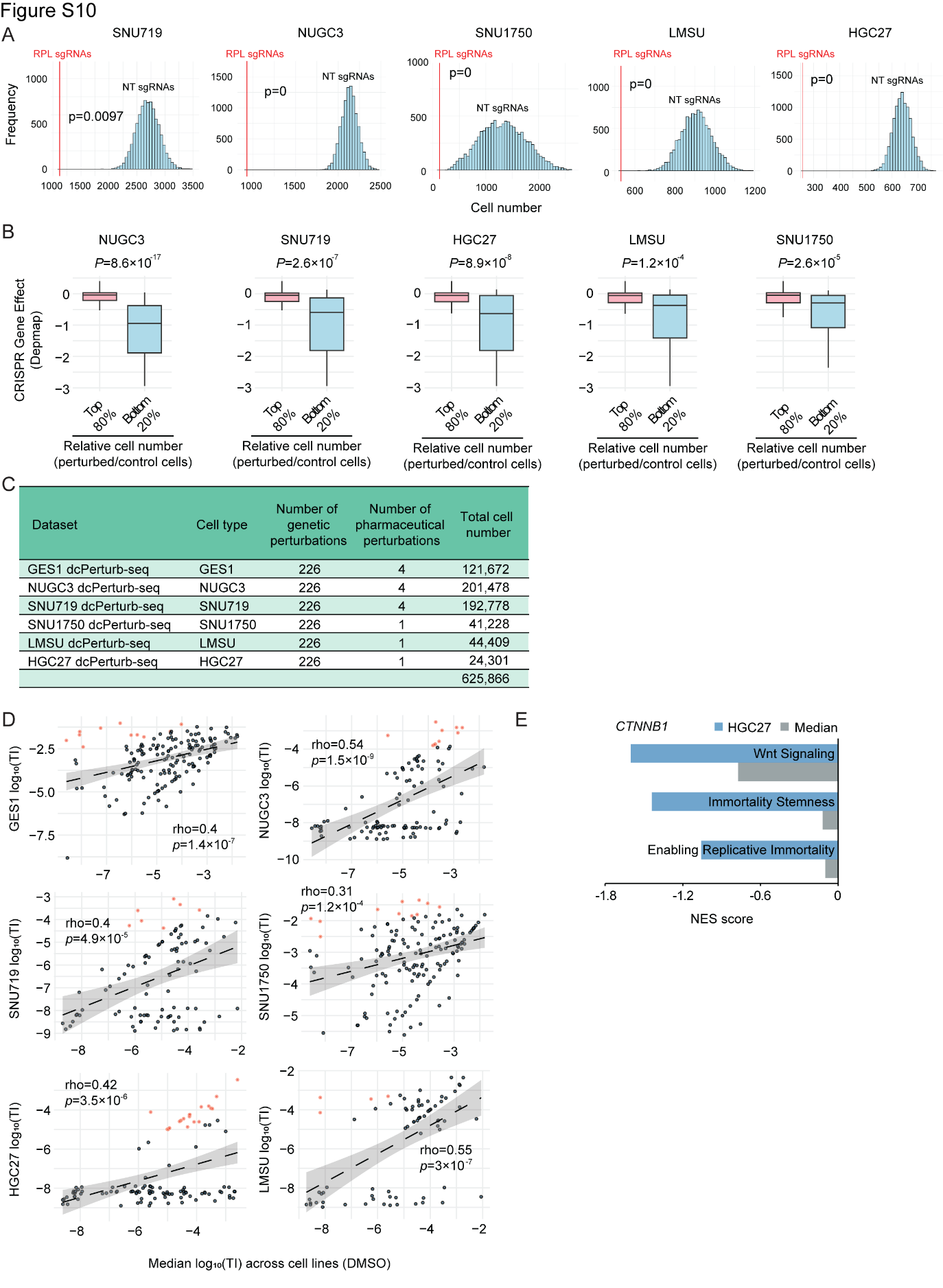

**Supplementary Figure 10 (Related to Figure 6)**

(A) Comparison of cell number distributions between sgRNAs targeting *RPL* genes and randomly selected sets of non-targeting sgRNAs. Significance was determined by an empirical permutation test.

(B) Boxplots comparing DepMap CRISPR gene effect for perturbations with bottom 20% and top 80% relative cell number (perturbed/control cells) (*P* values are adjusted p values calculated by Wilcoxon rank sum test).

(C) Summary of dcPerturb-seq dataset characteristics from this study.

(D) TI for gene perturbations in individual cell lines versus the median score across all lines under DMSO control, with outliers (orange) annotated. Significance was determined using Pearson correlations.

(E) Normalized enrichment scores (NES) for pathway analysis in *CTNNB1-*perturbed cells, compared to WT cells, for HGC27 and the median across six gastric cell lines under DMSO control. Normalized enrichment scores were generated from GSEA.

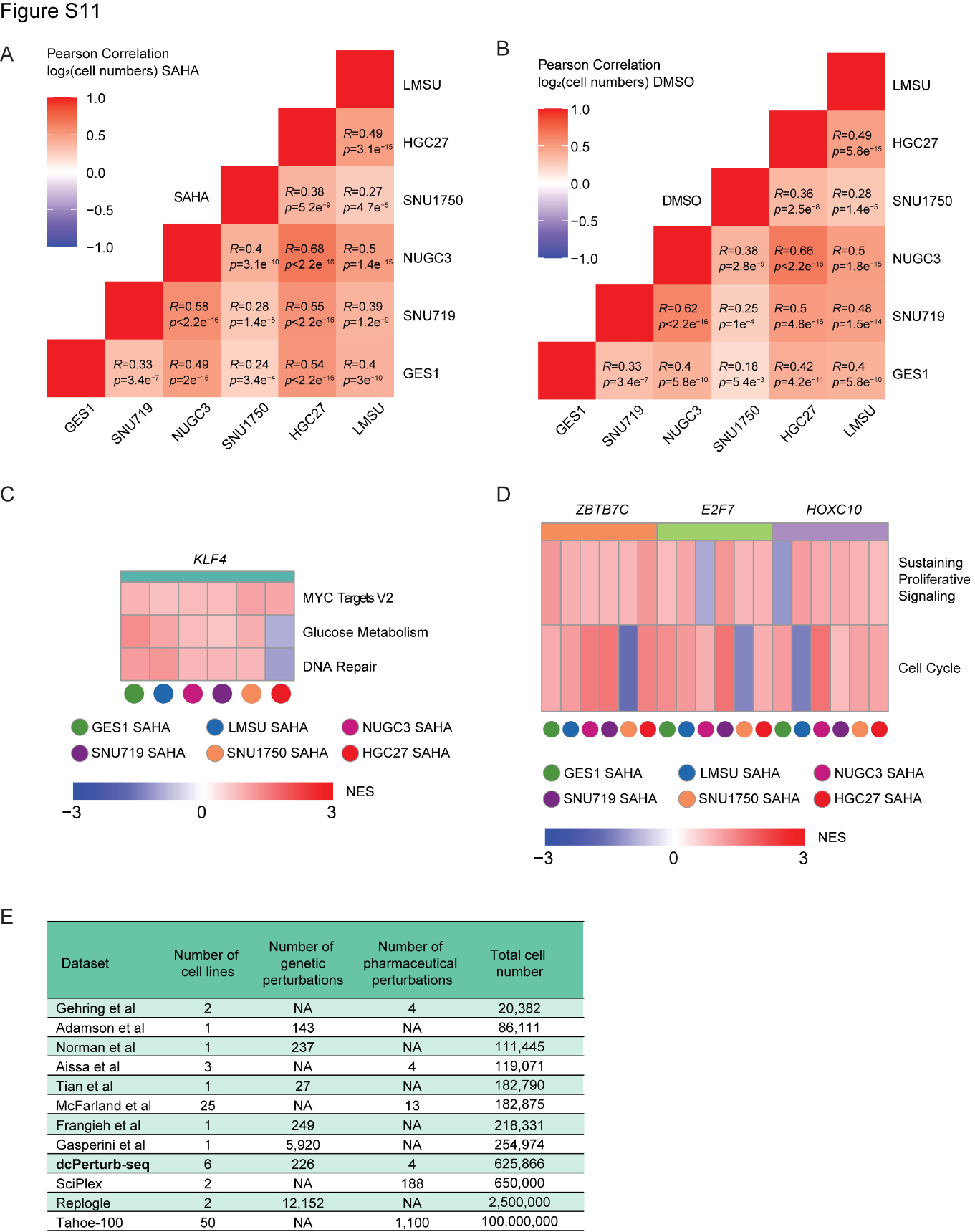

**Supplementary Figure 11 (Related to Figure 6 and Discussion)**

(A) Pairwise correlation of log_2_(cell number per gene) across cell lines under SAHA treatment. Significance was determined using Pearson correlations.

(B) Pairwise correlation of log_2_(cell number per gene) across cell lines under DMSO control condition. Significance was determined using Pearson correlations.

(C) Heatmap of normalized enrichment scores (NES) for pathway analysis in SAHA-treated *KLF4*-perturbed cells, compared to WT cells, across six gastric cell lines (GES1, LMSU, NUGC3, SNU719, SNU1750 and HGC27). Normalized enrichment scores were generated from GSEA.

(D) Heatmap of normalized enrichment scores (NES) for pathway analysis in SAHA-treated *ZBTB7C-*, *E2F7*- and *HOXC10*-perturbed cells, compared to WT cells, across six gastric cell lines (GES1, LMSU, NUGC3, SNU719, SNU1750 and HGC27). Normalized enrichment scores were generated from GSEA.

(E) Benchmarking of the current dcPerturb-seq dataset against comparable single-cell perturbation datasets (from scPerturb).

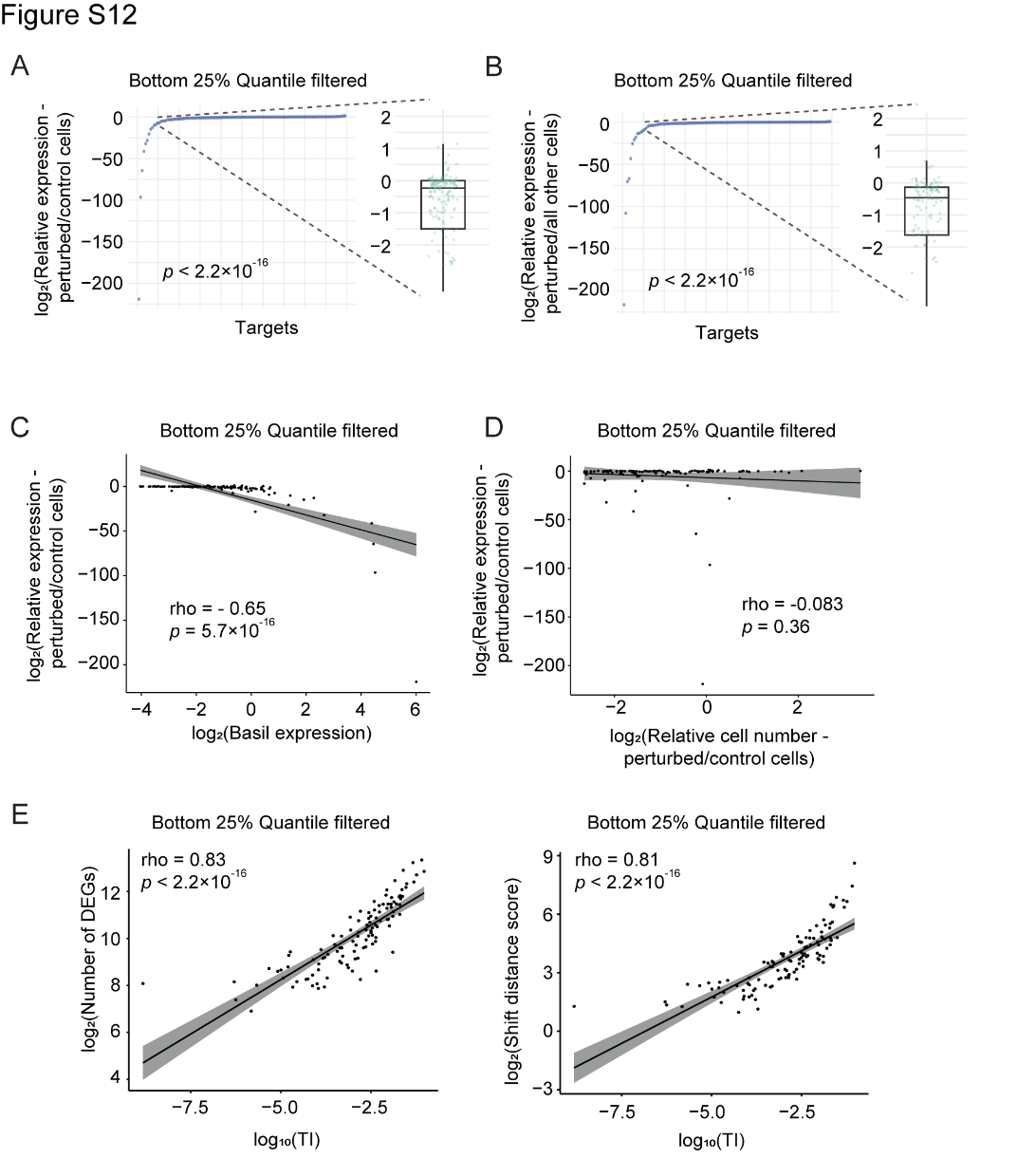

**Supplementary Figure 12 (Related to Methods)**

(A) Distribution of log_2_(relative target expression) comparing perturbed cells with all other cells for each target gene. Lowly expressed targets (bottom 25%) and those with low cell counts (bottom 25%) were filtered prior to analysis. Significance was determined using binomial tests.

(B) Distribution of log_2_(relative target expression) comparing perturbed cells with control cells for each target gene. Lowly expressed targets (bottom 25%) and those with low cell counts (bottom 25%) were filtered prior to analysis. Significance was determined using binomial tests.

(C) Correlation between basal expression and relative target expression (perturbed/control cells) for each target gene in GES1 cells. Lowly expressed targets (bottom 25%) and those with low cell counts (bottom 25%) were filtered prior to analysis. Significance was determined using Pearson correlations.

(D) Correlation between relative cell number (perturbed/control cells) and relative target expression (perturbed/control cells) for each target gene in GES1 cells. Lowly expressed targets (bottom 25%) and those with low cell counts (bottom 25%) were filtered prior to analysis. Significance was determined using Pearson correlations.

(E) Correlation between TI and the numbers of differentially expressed genes (DEGs) (left); Correlation between TI and the shift distance scores relative to wild-type (WT) cells (right) for each target gene in GES1 cells. Lowly expressed targets (bottom 25%) and those with low cell counts (bottom 25%) were filtered prior to analysis. Significance was determined using Pearson correlations.
